# A photoswitchable helical peptide with light-controllable interface / transmembrane topology in lipidic membranes

**DOI:** 10.1101/2021.03.10.434736

**Authors:** Mónica Gutiérrez-Salazar, Eduardo Santamaría-Aranda, Louise Schaar, Jesús Salgado, Diego Sampedro, Victor A. Lorenz-Fonfria

## Abstract

According to the three-step model, the spontaneous insertion and folding of helical transmembrane (TM) polypeptides into lipid bilayers is driven by three sequential equilibria: solution-to-membrane interface (MI) partition, unstructured-to-helical folding, and MI-to-TM helix insertion. However, understanding these three steps with molecular detail has been challenged by the lack of suitable experimental approaches to rapidly and reversibly perturb membrane-bound hydrophobic polypeptides out of equilibrium. Here, we report on a 24-residues-long hydrophobic α-helical polypeptide, covalently coupled to an azobenzene photoswitch (KCALP-azo), which displays a light-controllable TM/MI equilibrium in hydrated lipid bilayers. FTIR spectroscopy shows that dark-adapted KCALP-azo (*trans* azobenzene) folds as a TM α-helix, with its central TM region displaying an average tilt of 36 ± 4° with the membrane normal (TM topology). After *trans*-to-*cis* photoisomerization of the azobenzene moiety with UV light (reversed with blue light), spectral changes by FTIR spectroscopy indicate that the helical structure of KCALP-azo is maintained but the peptide experiences a more polar environment. Interestingly, pH changes induced similar spectral alterations in the helical peptide LAH_4_, with a well-characterized pH-dependent TM/MI equilibrium. Polarized experiments confirmed that the membrane topology of KCALP-azo is altered by light, with its helix tilt changing reversibly from 32 ± 5° (TM topology, blue light) to 79 ± 8° (MI topology, UV light). Further analysis indicates that, while the *trans* isomer of KCALP-azo is ~100% TM, the *cis* isomer exists in a ~90% TM and ~10% MI mixture. Strategies to further increase the perturbation of the TM/MI equilibrium with the light are briefly discussed.

## Introduction

Transmembrane (TM) α-helical proteins are involved in energy transduction and in the regulation of the traffic of molecules and information across biological membranes, processes of high biological relevance. In the cellular context, the insertion of TM segments of membrane proteins is in most cases assisted by a specialized protein complex called the translocon, and carried out almost synchronously with polypeptide synthesis by the ribosome.^1^ Nevertheless, mildly hydrophobic TM domains of C-terminally-anchored proteins can insert spontaneously into membranes, without translocon participation.^2^ Amphiphilic polypeptides, with antimicrobial, antitumoral, or apoptotic effects,^3^ provide another relevant example for the spontaneous membrane insertion of TM α-helices.

The well-established *three-step* model popularized by White and co-workers postulates that the spontaneous folding of TM helical protein fragments and peptides into lipid bilayers involves a solution-to-membrane interface (MI) partition, followed by folding into a helical structure, and completed by an MI-to-TM transition.^4^ This sequence of events has been broadly supported by experimental data^4^ and molecular dynamics simulations.^5^ However, experimental details on the partition, folding and insertion mechanisms, and their associated dynamics in both their forward and backward directions, remain poorly characterized. One reason for that stems from difficulties in perturbing TM helices out of their equilibrium. Hydrophobic peptides that form TM helices, although ideal models of membrane protein fragments, typically aggregate when added to aqueous solutions. Additionally, once inserted in membranes, they are very stable and unlikely to move back to the interface of the membrane, or to unfold.

One approach to experimentally study the insertion of peptides in lipidic membranes has been the use of pH-sensitive peptides, such as pHLIP, which exists in a water-soluble/MI equilibrium at neutral pH but inserts as a TM helix by decreasing the pH.^6^ This property was exploited to externally induce the binding, folding, and TM insertion of pHLIP *via* rapid changes in the medium pH by stopped-flow.^7,8^ However, the observed changes have been limited to a resolution of milliseconds and to single wavelength traces from Trp fluorescence spectroscopy, with limited information content. An alternative approach has been to induce the TM insertion of a membrane-bound peptide by rapidly increasing the membrane fluidity with a laser-induced temperature-jump (T-jump).^9^ Although this allowed for (sub)microsecond resolution and sample conditions compatible with infrared (IR) spectroscopy, the changes in the membrane fluidity lasted for less than 1 ms,^9^ making relevant millisecond and slower dynamical events inaccessible.

Some of the limitations of fast mixing methods^10^ and T-jumps^11^ for the folding/unfolding of water-soluble peptides have been avoided by their coupling to photoisomerizable organic molecules known as photoswitches.^12–14^ Most commonly, cysteine-reactive azobenzene derivatives have been linked to helical peptides, using as an anchor the thiol side chains of two cysteines.^15^ Changes in the azobenzene end-to-end length, typically from ~18 Å to ~10 Å as it photoisomerizes from *tran*s to *cis*,^16^ are used to distort/unfold the structure of the coupled helical peptide with light.^12,15,17^ The *trans* ismomer can be restored with light of a different wavelength, or thermally.^13^ By this approach, the folding/unfolding of helical soluble peptides has been studied by time-resolved IR difference spectroscopy.^14,18,19^ In addition, the coupling of photoswitches to soluble peptides has also been used to control their interaction with proteins or with DNA/RNA.^20,21^ In other cases, proteins themselves have been modified with photoswitches, allowing a direct optical control of their activity.^22–24^ In addition, photoswitches have been inserted in the molecular skeleton of many ligand compounds that are able to activate or inhibit proteins, making their affinity to target proteins light-sensitive, key for the development of the field of photopharmacology.^25^ There are also recent examples of the integration of photoswitches into cyclic β‑hairpin peptides that interact with membranes, derived from gramicidin S, with the goal of tuning their antimicrobial activity with light.^26–28^

In spite of the proven utility of photoswitches, the potential of using them on hydrophobic α-helical peptides remains unexplored. In this work, we synthesized the photoswitchable peptide KCALP-azo, derived from the KALP family of peptides, used in the past as models for monomeric TM α-helices, both in experiments and simulations.^29–31^ By means of FTIR absorption and difference spectroscopy we concluded that KCALP-azo folds in lipidic membranes as a TM α-helix in the dark (*trans* azobenzene conformation). Upon *trans*-to-*cis* azobenzene isomerization, induced with UV light, a fraction of KCALP-azo reorients to lay almost parallel to the membrane surface. This drastic change of the helix tilt, accompanied by an increase of the polarity sensed by the peptide backbone, informs about a change in the membrane topology of KCALP-azo from a TM to a MI state. This change was reversed by restoring the *trans* conformation of azobenzene with blue light illumination, demonstrating for the first time to our knowledge the reversible manipulation of a peptide TM/MI equilibrium with light.

## Results and discussion

We constructed a 24 residues-long hydrophobic peptide with cysteine residues at positions 7 and 18 (*i*, *i*+11) and sequence Ac-GKKLLACLLAALLAALLCALLKKA-NH_2_ (Fig. 1a), hereafter KCALP. Then, we incorporated the thiol-reactive cross-linker BCA (see Fig. 1c, left), *via* cysteine side chains, adapting a protocol initially derived to modify helical soluble peptides^32^ (see Experimental Section), giving the photoswitchable peptide KCALP-azo (Fig. 1b). As a spectroscopic control, we synthesized an azobenzene derivative that chemically mimics the product of the reaction of BCA with two thiol chains, named ThioAzo for short (Fig. 1c, right).

**Fig. 1.**
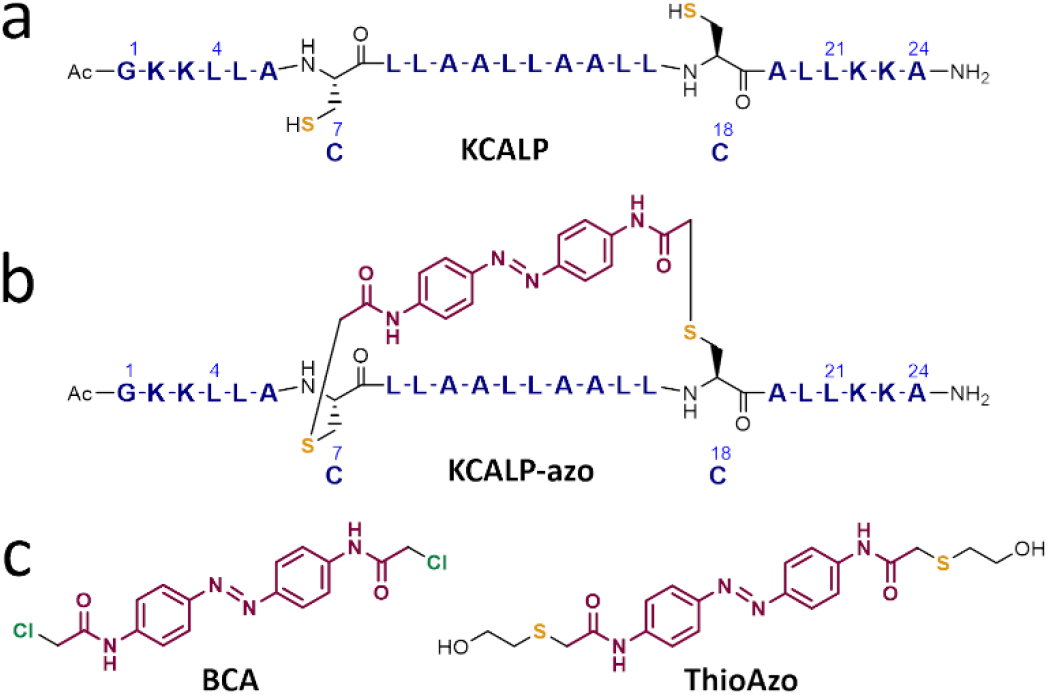
Structural comparison of KCALP, KCALP-azo, BCA and ThioAzo. (a) Primary structure of the peptide KCALP, with the chemical structure of Cys7 and Cys18 (C7 and C18). (b) Primary structure of the photoswitchable peptide KCALP-azo, with the chemical structure for Cys residues and for the azobenzene group. (c) Chemical structure of the cysteine-reactive azobenzene-based cross-linker BCA (left), and of the compound ThioAzo (right), vibrational model of the azobenzene group in KCALP-azo.

### Structural comparison of KCALP and KCALP-azo in dry POPC membranes

We gently dried POPC vesicles of reconstituted KCALP and KCALP-azo peptides, spontaneously forming oriented membranes (Fig. 2a). Unpolarized FTIR absorption spectra of these films are presented in Fig. 2b, which include the structure-sensitive backbone amide A (AA, *ν*N−H), amide I (AI, *ν*C=O), and amide II (AII, δN−H + *ν*C−N) vibrations,^33^ as well as *ν*CH_2_ and the ester *ν*C=O vibrations from the lipids.^34^ The amide A, I and II vibration frequencies at 3296 cm^−1^, 1659 cm^−1^ and 1546 cm^−1^, respectively, indicate that both KCALP and KCALP-azo adopt, predominantly, an α-helical structure.^33^ An amide I peak at a wavenumber as high as 1659 cm^−1^ has been only reported for TM helical peptides,^35^ while for soluble α-helical peptides the amide I peaks at 1650-1644 cm^−1^. ^36^

**Fig. 2.**
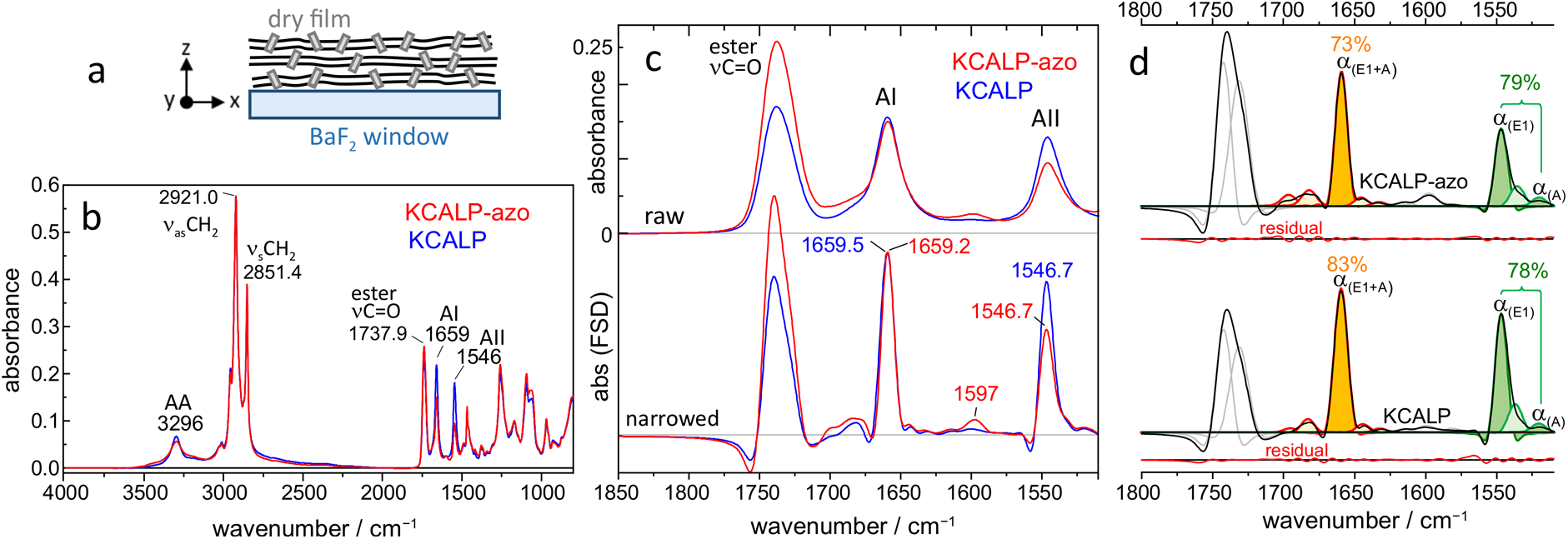
Structural comparison of KCALP and KCALP-azo in dried POPC membranes. (a) Sketch of oriented membranes. (b) Unpolarized FTIR spectra of KCALP-azo (red) and KCALP (blue). Peptide backbone amide A (AA), amide I (AI) and amide II (AII) vibrations and lipid *ν*CH_2_ and ester *ν*C=O vibrations are labelled. (c) Expanded 1850-1510 cm^−1^ region of the spectra in (b), before (top) and after (bottom) band-narrowing by FSD, scaled for the same amide I intensity. The band at 1597 cm-1, present only in KCALP-azo, originates from the *ν*C=C of azobenzene. (d) Band decomposition of FSD spectra using FSD-modified Voigtian bands (see Tables S1 and S2 for the band parameters). Bands in the 1700-1620 cm^−1^ region (red lines) and in the 1570-1515 cm^−1^ region (green lines) are assigned to amide I and amide II vibrations, respectively. Bands assigned to α-helices are colour-filled (orange for amide I and olive for amide II bands), and their area percentages are indicated. Fitting residuals

Figure 2c (top) expands the region between 1850 and 1510 cm^−1^, which includes the amide I and II vibrations. To resolve more details, we applied Fourier self-deconvolution, FSD, a mathematical tool for band-narrowing (Fig. 2c, bottom).^37^ Note the high spectral similitude between KCALP-azo and KCALP for the amide I band after FSD, an indication that the photoswitch perturbs minimally the original structure of KCALP, as expected for an α-helical peptide with an azobenzene unit linked to *i* and *i*+11 cysteine residues.^15^ The symmetric and rather narrow amide I and II bands from KCALP-azo and KCALP contrast with those from α-helix-rich membrane proteins (Fig. S1), supporting a monomeric and homogenous structure for the two peptides. Incidentally, the band at 1597 cm^−1^ in KCALP-azo (see Fig. 2c), can be assigned to *ν*C=C vibrations of the azobenzene group.^38^

To obtain quantitative information about the secondary structure of KCALP and KCALP-azo, we decomposed their amide I and II envelops into sums of subcomponent bands by nonlinear least-squares. This decomposition is more robust when conducted on FSD spectra, in particular when using FSD-modified Gaussian, Lorentzian or Voigtian bands, instead of their conventional versions.^39,40^ Figure 2d shows the band decomposition using FSD-modified Voigtian bands, and Tables S1 and S2 collect the estimated parameters for each fitted Voigtian band.^39,40^ For an α-helix, amide vibrations couple into three vibrational modes, two of which are IR active: A and E_1_.^33^ Because the A and E_1_ modes strongly overlap for amide I vibrations, with a splitting of only ~2-7 cm^−1, 33,41^ they are observed as a single band at ~1659 cm^−1^ for both KCALP and KCALP-azo (Fig. 2d). For the amide II vibration the splitting of the E_1_ and A modes is 20-30 cm^−1, 33,36,41^ large enough to be well-resolved into two bands at ~1547 (E_1_) and ~1520 (A) cm^−1^ (Fig. 2d). The total band area assigned to helical structures in KCALP (colour-filled bands in Fig. 2d) corresponded to 83% and 78% of the total amide I and amide II area, respectively (Fig. 2d, bottom). For KCALP-azo, on the other hand, these percentages were 73% and 79% (Fig. 2d, top).

A more accurate estimation of the helical content of KCALP and KCALP-azo require us to take into account other groups with vibrations contributing to the amide I-II region. These include the NH_3_+ group from Lys (at ~1630 and ~1525 cm^−1^ in H_2_O),42 the secondary amide from the acetylated N-termini (at ~1630 cm^−1^ and ~1580 cm^−1^ in H_2_O),^43^ and the primary amide from the amidated C-termini (at ~1670 cm^−1^ and ~1610 cm^−1^ in H_2_O).^42^ KCALP-azo contains two additional secondary amide groups located at the azobenzene linker (see Fig. 1b), with expected bands at ~1700-1680 cm^−1^ and at ~1530-1500 cm^−1^ in aprotic organic solvents of low/medium polarity.^44^ Upon considering the expected wavenumber and absorption coefficient of these vibrations relative to those of the amide backbone group,^42^ we concluded that both peptides are roughly 85% α-helical.

In spite of the high resemblance between the vibrational spectra of KCALP and KCALP-azo, a good proxy for structural similarity, KCALP-azo displays an amide I / amide II intensity ratio notably higher than KCALP (Fig. 2c). This observation suggests possible differences in their helix tilt with respect to the lipid membrane normal. To clarify this point, we conducted polarized-light FTIR experiments, presented in Fig. S2, from where we estimated the average helix tilt, ⟨*β*⟩, to be 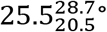 for KCALP and 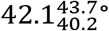 for KCALP-azo. Thus, the coupling of azobenzene to KCALP does not perturb its helical structure nor its TM topology, albeit increases its average helix tilt by ~16.5°. Note that here, and throughout the text, we provide the most likely average tilt angle with an asymmetrical 96% confidence level, but in some cases we used for simplicity symmetrical confidence limits, *e.g.*, 24.5 ± 4° instead of 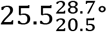.

### Structure of KCALP-azo in hydrated POPC membranes

Films of KCALP-azo in POPC membranes were hydrated for all subsequent studies, either with ~450 (film A, Fig. 3b) or with ~950 water molecules per peptide (film B, Fig. S3). In either case, the lipid *ν*_s_CH_2_ vibration, at ~2853.4 cm^−1^ (insets in Fig. 3b and Fig. S3), confirmed that the membranes were in the fluid phase.^34^ We digitally subtracted the background absorbance from liquid water, revealing the structure-sensitive amide I and amide II vibrations of the peptide backbone (Fig. 3b and Fig. S3, blue trace). After FSD, the amide I and II bands of KCALP-azo peaked at 1656.8 ± 0.2 cm^−1^ and 1548.1 ± 0.2 cm^−1^ (Fig. S4a).

**Fig. 3.**
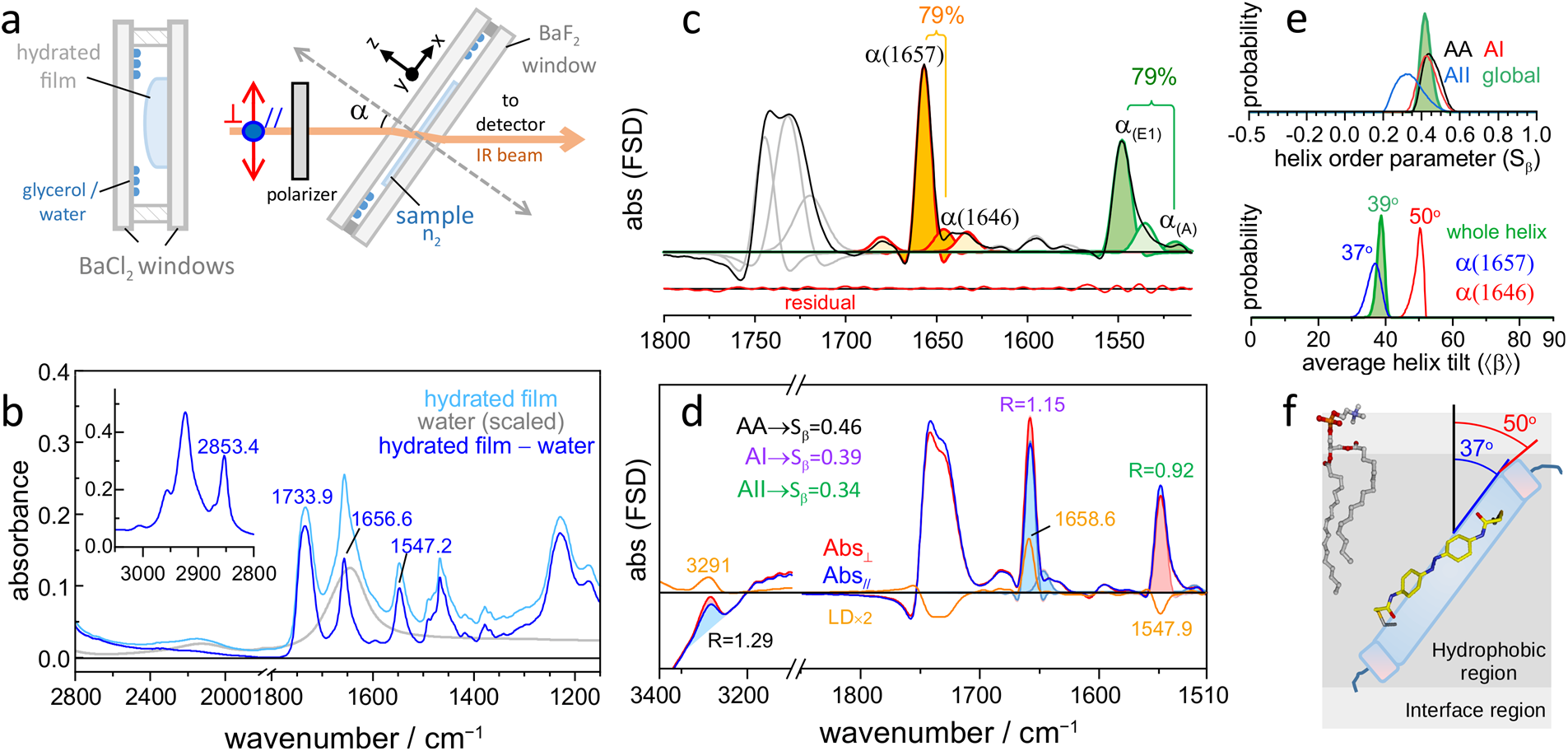
Structure and orientation of KCALP-azo in hydrated POPC membranes. (a) Schematic experimental setup. (b) Unpolarized FTIR spectra of KCALP-azo before (light blue trace) and after (blue trace) subtraction of the water absorbance (grey trace). (c) Band decomposition of the FSD spectrum of KCALP-azo (see Table S3). (d) FSD polarized FTIR spectra of KCALP-azo (α = 50°). Dichroic ratios (*R*) from helices are given for each amide vibration, together with the second order parameter of the helix tilt (*S*_*β*_). The LD spectrum is shown in orange, two times enlarged for clarity. (e, top) Probability distributions of *S*_*β*_ from each amide vibration (amide A, black; amide I, red; amide II, blue), and its global estimate (green, colour-filled). (e, bottom) Probability distributions of the average helix tilt for: the whole helix (green), and the helix segments corresponding to the bands at 1657 cm^−1^ (blue) and 1645 cm^−1^ (red). (f) Schematic model of KCALP-azo in a hydrated POPC membrane. The peptide helical structures are represented with cylinders. The hydrophobic core and interface regions of the lipid membrane is shown in grey and light grey, respectively, with a single POPC molecule depicted for illustration purposes.

To obtain a secondary structure estimate we focused on film A, because its lower water content made spectral corrections and the subsequent band-decomposition more reliable. The main amide I component band, with 66% of the area, is located at 1656.8 cm^−1^ (Fig. 3c and Table S3), in the range for TM α-helices in proteins and peptides.^35,45^ We assigned the band at 1646.0 cm^−1^, with 13% of the area (Fig. 3c and Table S3), to the amide I of hydrated α-helices.^46,47^ The assignment of amide I bands at 1656.8 and 1646.0 cm^−1^ to helical structures is consistent with the band-decomposition of the amide II, giving a 79% of helical structures (Fig. 3c and Table S3). We obtained comparable results for the hydrated film B (Table S4). Considering potential overlapping spectral contributions, discussed in the previous section, the estimated percentage of helical structures was ~85%. Therefore, the helicity of KCALP-azo does not significantly change upon hydration, although a portion of the TM helix, likely at the vicinity of the membrane interface, becomes involved in H-bonds with interfacial water molecules.

### Orientation of KCALP-azo in hydrated POPC membranes

We recorded FTIR absorption spectra using IR light with a polarization parallel (Abs_⁄⁄_) or perpendicular (Abs_⊥_) to the axis of rotation of the sample window (Fig. 3a). The resulting spectra were water-corrected and band-narrowed by FSD (Fig. 3d). The linear dichroism spectrum (LD = Abs_⊥_ − Abs_⁄⁄_) shows a positive amide A, a positive amide I, and a negative amide II band (Fig. 3d, orange), similarly to KCALP-azo in dry POPC membranes (Fig. S2b). This sign-pattern is characteristic for TM helical peptides^48^ and proteins.^49–51^ In contrast, the pH-sensitive helical peptide LAH_4_, adopting a MI topology at low pH,52 displays the opposed sign-pattern, as shown in Fig. S2b (bottom).

To estimate the average helix tilt of KCALP-azo with respect to the membrane normal, ⟨*β*⟩, we first determined dichroic ratios, R, as the Abs_⊥_/ Abs_⁄⁄_ area ratio of amide vibrations from helical structures (Fig. 3d). Then, we estimated the second order parameter of the helix tilt, defined as *S*_*β*_ = (3⟨cos^2^*β*⟩ −- 1)⁄2 (the brackets represent space and time-averaged values), using the relation:^53,54^

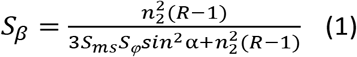

In this equation *n*_2_ represents the refractive index of the hydrated film, *S*_*ms*_ its mosaic spread (*i.e.*, the second order parameter of the lipid membrane normal with respect to the normal of the solid support), and *S*_*φ*_ the second order parameter of the angle between the transition dipole moment of an amide vibration and the helix axis. From the *R* values for the three amide vibrations (Fig. 3d), and by considering the most likely values of the parameters involved in Eq. 1 as well as their uncertainty (see Experimental Section), we obtained three probability distributions for *S*_*β*_ (Fig. 3e-top, black, red and blue traces). Combining them, we obtained a global estimate of 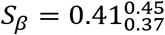 (Fig. 3e, top, green trace). From the definition of *S*_*β*_ (see above), and applying the approximation ⟨*cos*^2^*β*⟩ ≈ *cos*^2^⟨*β*⟩ (whose suitability is demonstrated below), we estimated the average helix tilt of KCALP-azo to be 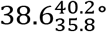 (Fig. 3e, bottom, green trace).

KCALP-azo contains at least two distinguishable types of α-helices, resolved as two distinct amide I bands at ~1657 and ~1646 cm^−1^ (Fig. 3c and Table S3). We were able to determine their individual dichroic ratios from the band-decomposition shown in Fig. 3d (*R* = 1.17 and 1.08, respectively) and, thus, their average tilts (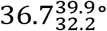 and 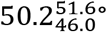, respectively), as shown in Fig. 3e (bottom, blue and red trace). Considering their wavenumbers (~1657 and ~1646 cm^−1^), their average tilts (~37° and ~50°) and their areas (~70% and ~15%), we assigned the first of the two bands to the core segment of the TM-helix, and the second band to shorter helical segments at the peptide ends, near the interfacial region of the membrane, schematically illustrated in Fig. 3f.

As we have noted, we rely on the approximation ⟨cos^2^*β*⟩ ≈ *cos*^2^⟨*β*⟩ to obtain average helix tilt values, ⟨*β*⟩, from polarized IR experiments, *i.e.*, we need to assume that *S*_*β*_ ≈ (3cos^2^⟨*β*⟩ − 1)⁄2. This approximation is only exact when the tilt angle takes a discrete value, which is not the case for helical peptides in membranes.^55,56^ To evaluate the error made, we computed exact values for *S*_*β*_ and ⟨*β*⟩ for a family of distributions of tilt angles (Fig. S5a,b), and used the approximation *S*_*β*_ ≈ (3*cos*^2^⟨*β*⟩ − 1)⁄2 to determine back ⟨*β*⟩ from *S*_*β*_. The discrepancy between exact and the recovered values for ⟨*β*⟩ was < 2.5° (in most cases < 1°, Fig. S5b), smaller than our statistical error, validating our approach.

### Photoisomerization of azobenzene in KCALP-azo

Once we determined that in hydrated POPC membranes ~85% of the backbone of KCALP-azo folds as α-helix, the core of which (~70%) forms a TM hydrophobic segment with an average tilt of 36 ± 4°, we focused on studying how photoisomerization of the covalently bound azobenzene group perturbs the structure and orientation of KCALP-azo in hydrated POPC membranes. But before that, we needed to quantify to which degree we could photoisomerize the azobenzene moiety of KCALP-azo under our experimental conditions.

The UV-Vis absorption spectrum of KCALP-azo in hydrated POPC membranes (Fig. 4a, red trace) shows a strong absorption band at 368 nm, characteristic for the electronic π→π* transition of azobenzene in *trans* configuration.^21^ After UV illumination for 2 seconds (λ_max_ = 365 nm, 400 mW/cm^2^) the absorption at 368 nm notably decreases, while the one at ~454 nm increases slightly (Fig. 4a, blue trace), indicating the partial formation of the *cis* conformer of azobenzene.^21^ Blue illumination for 2 seconds (λ_max_ = 447 nm, 200 mW/cm2) largely recovers the initial absorption spectrum (Fig. 4a, green trace). A UV-Vis difference spectrum between the *trans* and *cis* conformers is shown in Fig. 4b. Note that experiments made in parallel by FTIR difference spectroscopy confirmed that 2 s of illumination was sufficient to reach a photostationary state, PSS (Fig. S6).

**Fig. 4.**
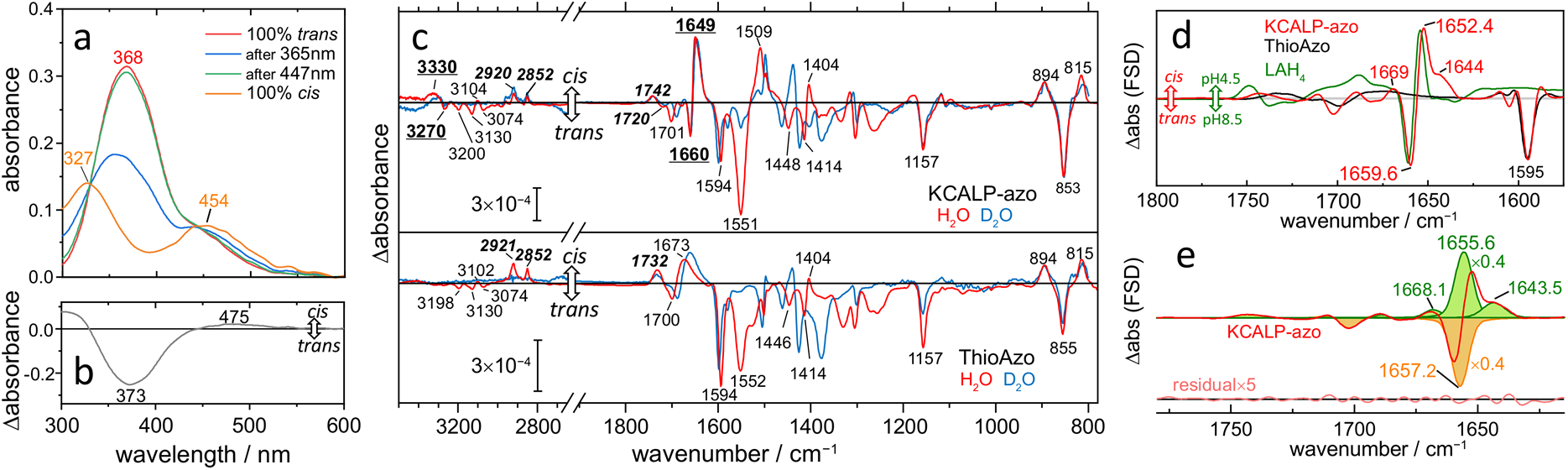
Light-induced changes after photoisomerization of KCALP-azo in hydrated POPC membranes. (a) UV-Vis absorption spectrum of dark-adapted KCALP-azo (100% *trans* conformation), after 365 nm (blue line), and after 447 nm (green line) illumination. The spectrum for 100% *cis* KCALP-azo (orange line) is an estimate. (b) UV-Vis difference spectrum between *cis* and *trans* KCALP-azo. (c, top) Light-induced (365 nm-*minus*-447 nm) unpolarized FTIR difference spectra of KCALP-azo in membranes hydrated with H_2_O (red trace) and D_2_O (blue trace). The bands are labelled following their tentative assignment: peptide backbone (bold and underlined), lipid (italics), and photoswitch (plain). (c, bottom) Light-induced FTIR difference spectra of ThioAzo in hydrated POPC membranes. (d) Light-induced FTIR difference spectra of KCALP-azo (red) and ThioAzo (black), together with a scaled pH-induced FTIR difference spectrum of LAH_4_ (green), all band-narrowed with FSD. (e) Band-fitting of the FTIR difference spectrum of KCALP-azo in (d), with band parameters in Table S5. Note that the fitted bands at 1657.2 and 1655.6 cm^−1^ are scaled by 0.4 to keep their intensity within the displaying limits.

Although we could not obtain 100% of *cis* KCALP-azo in POPC membranes by illumination, we could estimate its spectrum (Fig. 4a, orange trace). For that, we took as guidance the spectrum of 100% *cis* KCALP-azo measured in acetonitrile/water after its isolation by HPLC (Fig. S7c). Then, considering that dark-adapted KCALP-azo is 100% *trans*, we estimated a 53% *cis* / 47% *trans* PSS after 365 nm illumination, and a 4% *cis* / 96% *trans* PSS after 447 nm illumination in POPC membranes; *i.e.*, we could achieve a maximum of ~50% conversion between the *trans* and *cis* forms by alternating UV and blue light illumination. The above estimates are consistent with PSS values determined for KCALP-azo in acetonitrile/water solution by HPLC: 2% *cis* / 98% *trans* in the dark-adapted state, 48% *cis* / 52% *trans* after 365 nm illumination, and 13% *cis* / 87% *trans* after 447 nm illumination (Fig. S7b).

### Light-induced structural changes of KCALP-azo

UV-Vis spectroscopy confirmed the successful photoisomerization of the azobenzene group of KCALP-azo in hydrated POPC membranes, but it provided no clue about the accompanying structural changes in the peptidic part. Thus, we performed parallel experiments on the hydrated film B of KCALP-azo, measuring a FTIR difference spectrum after 2 s of UV illumination. The resulting spectrum was noisy, affected by a baseline drift, and included spectral signatures of heating (Fig. S8b). To improve the spectral quality and to reduce heating artefacts, we alternated short light pulses of 365 nm (100 ms) and 447 nm (200 ms), repeating this cycle ~500 times, and averaging the result (see Fig. S8a for a data acquisition scheme). The improvement in the spectral quality was remarkable (Fig. S8b). As a drawback, the intensity of the light-induced FTIR difference spectrum of KCALP-azo decreased by a factor of ~2.5 (Fig. S8b, red trace), *i.e.*, a reduction of the photoconversion between the *trans* and *cis* forms of KCALP-azo from a ~50% to a ~20%. This difference FTIR spectrum is reproduced in Figure 4c (top, red trace). A similar light-induced FTIR difference spectrum was obtained for the hydrated film A (see Fig. S4b).

Before interpreting the light-induced vibrational changes in KCALP-azo (Fig. 4c, top), it was necessary to distinguish bands which originated from vibrations localized in the peptidic part from those coming from the azobenzene group. To achieve that, without resorting to complex and expensive isotope labelling, we performed equivalent experiments on ThioAzo in POPC membranes (Fig. 4c, bottom), used as a vibrational model of the azobenzene photoswitch in KCALP-azo (see Fig. 1). Most of the bands in the light-induced FTIR difference spectrum of KCALP-azo are also present in ThioAzo, *i.e.*, they are assignable to vibrations localized in the azobenzene photoswitch. Among them, the bands at 1700 (−) and 1673 (+) cm^−1^ in ThioAzo and the band at 1701 (−) cm^−1^ in KCALP-azo (Fig. 4c) can be assigned to amide I vibrations of the amide group at the photoswitch linker (see Fig. 1), while the band at 1552 (−) cm^−1^ in ThioAzo and at 1551 (−) cm^−1^ in KCALP-azo (Fig. 4c) likely correspond to amide II vibrations. The former bands downshifted by 10-13 cm^−1^ and the latter virtually vanished upon hydration with D_2_O (Fig. 4c, blue), as expected from their assignment.^44^ Given the known requirements for hydrogen/deuterium exchange (HDX),^57^ we can conclude that the two amide N−H groups at the photoswitch linker are largely free from H-bonding and, at least transiently, accessible to solvent water molecules.

Two intense bands at 1660 (−) and 1649 (+) cm^−1^ are observed only in the light-induced FTIR spectrum of KCALP-azo (Fig. 4c, top), which can be assigned to changes in backbone amide I vibrations between the *trans* (unperturbed) and *cis* (perturbed) conformations of KCALP-azo. Likewise, two bands at ~3330 (+) / ~3270 (−) cm^−1^ in KCALP-azo, absent in ThioAzo, can be assigned to changes in amide A vibrations of the peptide backbone. Very importantly, these backbone amide bands did not show up upon illumination of membranes containing equimolar quantities of KCALP and ThioAzo (Fig. S9), meaning that illumination changed the structure of KCALP only when azobenzene was covalently bound to the peptide. Upon incubation of lipid-reconstituted KCALP-azo in D_2_O, the backbone amide I bands downshifted by less than 2.5 cm^−1^ in the light-induced FTIR difference spectrum (Fig. 4c top, compare red and blue traces), which contrasts with the expected 12 cm^−1^ downshift for a complete HDX.^44^ Therefore, most of the backbone amide N−H groups of KCALP-azo affected by illumination are HDX-resistant and, thus, involved in stable intramolecular H-bonding in both the unperturbed (*trans*) and perturbed (*cis*) states. Because D_2_O removed amide II bands from the photoswitch, but not from the peptide backbone, we could access changes from previously hidden backbone amide II vibrations (Fig. 4c, blue traces). Only a small negative amide II band at ~1550 cm^−1^ was observed, without any accompanying positive band (Fig. 4c top, blue), indicating that the amide II vibration is largely insensitive to the structural changes caused by photoisomerization of KCALP-azo.

Although the 11 cm^−1^ downshift of the backbone amide I, from 1660 (−) to 1649 (+) cm^−1^, might suggest a notable perturbation of the helical structure of KCALP-azo upon azobenzene photoisomerization, we should be aware that the apparent separation of bands with an opposed sign is always limited by their bandwidth.^58^ Indeed, after band narrowing by FSD, which reduces bandwidths, the same two bands were resolved only ~7 cm^−1^ apart, at 1659.6 (−) and 1652.4 (+) cm^−1^ (Fig. 4d). To resolve the bands contributing to the FTIR difference spectrum fully free from overlap effects, we relayed on band decomposition by nonlinear least-squares fitting. After this procedure, the two main amide I bands were resolved at 1657.2 (−) and 1655.6 (+) cm^−1^, only 1.6 cm^−1^ apart (see Fig. 4e and Table S5). The band-decomposition also revealed that both the position and width of the negative amide I band are very close to those of the main amide I band in the absorbance spectrum of KCALP-azo (Table S5 *vs* Table S4): 1657.2 *vs* 1657.0 cm^−1^ and 14.2 vs 16.6 cm^−1^, respectively. This notable similitude favours the idea that the light-induced changes affect the whole hydrophobic segment of the TM helix of KCALP-azo.

To interpret the 1.6 cm^−1^ downshift of the amide I of KCALP-azo caused by light, we briefly moved our attention to the pH-sensitive helical peptide LAH_4_. A dry film of POPC-reconstituted LAH_4_ at pH 8.5, where the peptide is known to fully adopt a TM topology,^52,59^ displayed a main amide I band at 1658.1 cm^−1^ (Fig. S10). The amide I maximum downshifted by 1.9 cm^−1^, to 1656.2 cm^−1^, at pH 4.5 (Fig. S10), a condition where the peptide fully adopts a MI topology.^52,59^ The amide I frequency downshift can be explained by an increase of the medium polarity as the peptide moves to the membrane interface at low pH.60 The amide II band of LAH_4_ was largely insensitive to the change in membrane topology (Fig. S10). In summary, the spectral differences associated with a change in membrane topology of LAH_4_ show a remarkable resemblance to those observed for KCALP-azo (compare red and green traces in Fig. 4d), suggesting that KCALP-azo might experience a light-induced change in its membrane topology.

Although most of the amide I band at 1657.2 cm^−1^ shifts to 1655.6 cm^−1^ upon azobenzene photoisomerization, part of it also shifts to 1643.5 and 1668.1 cm^−1^ (Fig. 4d,e and Table S5). The positive amide I band at 1643.5 cm^−1^ is akin to the band at 1645.6 cm^−1^ in the FTIR absorption spectrum of KCALP-azo (Fig. 3c and Table S4), indication that part of the TM helical segment might establish H-bonds with water after adopting a MI topology. On the other hand, the positive amide I band at 1668.1 cm^−1^ might originate from weakly H-bonded backbone amide carbonyl groups,^44^ formed as a result of structural distortions in the TM helical segment of KCALP-azo.

### Light-induced changes in the helix tilt of KCALP-azo

In order to detect changes in the tilt of the helix of KCALP-azo, we first performed light-induced polarized FTIR difference spectroscopy in combination with attenuated total reflection (ATR). We conducted these experiments on oriented films hydrated with D_2_O, providing access to amide A, I, and II bands from the peptide. Changes in the helix tilt can be easily gauged by transforming polarized ATR-FTIR difference spectra (ΔAbs_⊥_and ΔAbs_⁄⁄_) into difference spectra in the plane (ΔAbs_*x-y*_) and perpendicular (ΔAbs_*z*_) to the surface of the ATR crystal (see Fig. 5a, Inset).^61^ An isotropic difference spectrum, ΔAbs_*iso*_, free from orientation effects, is obtained as: ΔAbs_*iso*_ = (ΔAbs_*z*_ + 2×ΔAbs_*x-y*_)/3.^61^ To explain how ΔAbs_*x-y*_ and ΔAbs_*z*_ help to detect changes in the tilt of helices, Figure 5a includes cartoons of a simple idealized helix changing its tilt from 0° to 90° with respect to the *z* axis. When the transition dipole moment aligns close to the direction of the helix axis, like for the amide A (φ = 27-33°),^41,62,63^ a larger helix tilt causes the absorption to decrease in the *z* direction and to increase in the *x-y* plane, leading to a negative band in the ΔAbs_*z*_ spectrum and a positive band in the ΔAbs_*x-y*_ spectrum.^61^ This was indeed the observed pattern for the amide A band of KCALP-azo (Fig. 5a, green shaded area), compelling evidence for an increased helix tilt upon *trans*-to-*cis* azobenzene photoisomerization. Being the transition dipole moment of the amide II close to perpendicular to the helix axis (φ = 70-76°),^41,62,63^ a larger helix tilt should rise the absorption in the *z* direction while decreasing it in the *x-y* plane. Indeed, we observe a positive amide II band in the ΔAbs_*z*_ spectrum and a negative one in the ΔAbs_*x-y*_ spectrum (Fig. 5a, orange shaded area), further confirming a light-induced increase of the helical tilt of KCALP-azo.

**Fig. 5.**
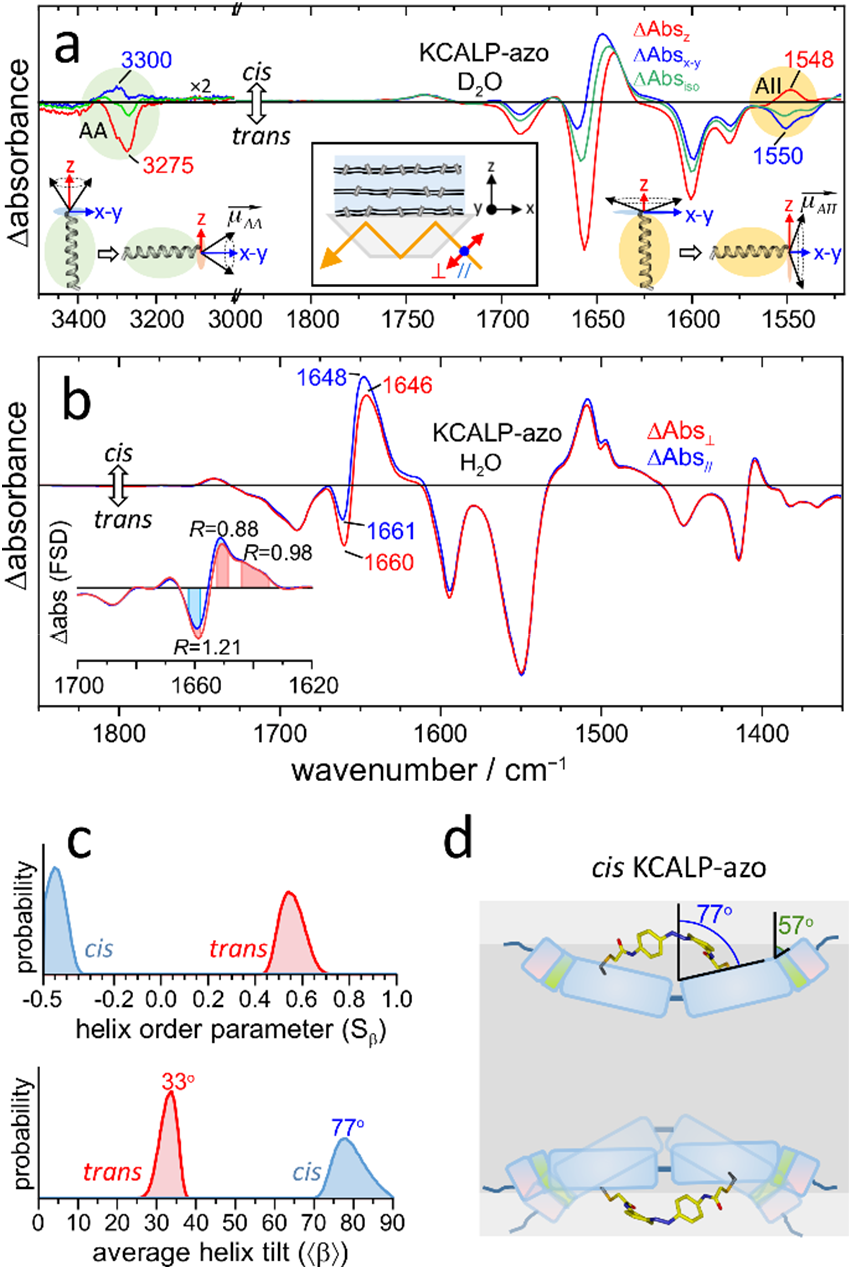
Light-induced orientation changes of the α-helix of KCALP-azo in hydrated POPC membranes. (a) Light-induced FTIR difference spectra in D_2_O in the plane (ΔAbs_*x-y*_, red) and perpendicular to the sample surface (ΔAbs_*z*_, blue), obtained by ATR with polarized light (see inset). ΔAbs_*iso*_ (green) is the isotropic difference spectrum. Changes in ΔAbs_*x-y*_ and ΔAbs_*z*_ for the amide A (green-shared area) and amide II (orange-shaded area) bands indicate an increase of the helix tilt of KCALP-azo upon *trans*-to-*cis* photoisomerization. For clarity, the tilt increase is illustrated with cartoons showing a 0° to 90° change. (b) Light-induced polarized FTIR difference spectra in H_2_O, measured by transmission (α = 50°). The inset shows the amide I region after FSD. The dichroic ratio, *R*, for two positive bands (*cis* state) and one negative band (*trans* state) is indicated. (c) Probability distributions for the second order parameter (top) and for the average tilt angle (bottom) of the main helix segment of KCALP-azo in *trans* (red) and *cis* (blue) conformations. (d) Schematic structural model of *cis* KCALP-azo. Helical

Only the amide I frequency of the helix changed sufficiently with the tilt as to resolve both positive (*cis*) and negative (*trans*) bands in both ΔAbs_*z*_ and ΔAbs_*x-y*_. Because the negative band was more intense in ΔAbs_*z*_ than in ΔAbs_*x-y*_, but the other way around for the positive band, we can conclude that the helix tilt of KCALP-azo is smaller than the magic angle (54.7°) after blue illumination (*trans* azobenzene) but larger after UV illumination (*cis* azobenzene). To obtain quantitative values for the helix tilt we conducted additional light-induced polarized FTIR difference experiments, but this time by transmission and with samples hydrated with H_2_O (Fig. 5b). We determined the dichroic ratio of the negative and positive amide I bands by integrating their area after FSD (Fig. 5b, Inset). For the negative band *R* was 1.21, and we arrived at 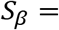 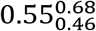 and at 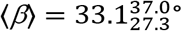 (Fig. 5c, red). This last estimate is in statistical agreement with the average helix tilt determined for the hydrophobic TM helical segment of KCALP-azo in the dark, 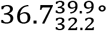 (Fig. 3e, bottom, blue trace), and indication that this is the segment of KCALP-azo whose tilt is perturbed by light.

From the dichroic ratio of the positive band, 0.89 (Fig. 5b, inset), we arrived at 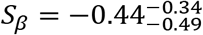 and at 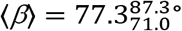 for *cis* KCALP-azo (Fig. 5c, blue). Figure S5d displays several distributions of *β* compatible with *S*_*β*_ = −0.44, spanning from a perfectly homogenous helix tilt of *β* ≈ 77° to a broad Gaussian distribution of *β* angles centred at ~90°. Note that a helix tilt ≥77° for a 24 residues-long peptide appears incompatible with a TM topology, even if considering possible snorkelling effects of the Lys charged residues in the helix termini. Instead, this tilt agrees well with those previously described for MI helical peptides.^64–67^ Incidentally, the positive band at 1643.5 cm^−1^ (Fig. 4e), assigned to hydrated helical segments (see above), displayed a dichroic ratio of 0.98 (Fig. 5b, inset), from where we estimated that its average helix tilt is 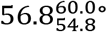.

Figure 5d shows two structural models for the MI state of *cis* KCALP-azo compatible with our experimental results. The first model is a V-shaped helical conformation with a static orientation (Fig. 5d, top). The second model the helix exhibits a dynamic orientation, fluctuating between a flat alignment at the membrane interface and a V-shaped conformation (Fig. 5d, bottom). Both models can explain the average tilt of the hydrophobic helix segment of *cis* KCALP-azo (~77°). Although we lack direct information about the peptide azimuthal rotation, we tentatively placed it facing the membrane interface region given that the polarity of azobenzene markedly increases in its *cis* isomeric state.^22^

### Fraction of KCALP-azo moving to the membrane interface after azobenzene photoisomerization

We reconstructed the FTIR absorption spectrum of pure *cis* KCALP-azo by taking into account that under our illumination conditions the change in the isomeric state of azobenzene occurs only for ~20% of the KCALP-azo molecules. This was done by simply adding to the FTIR absorption spectrum of dark-adapted KCALP-azo (100% *trans*), the light-induced FTIR difference spectrum multiplied by 5. The resulting 100% *cis* spectrum, presented in Fig. 6a after FSD, displays the expected reduction in the intensity of the 1596 cm^−1^ band from the *ν*C=C of azobenzene.^68^

**Fig. 6.**
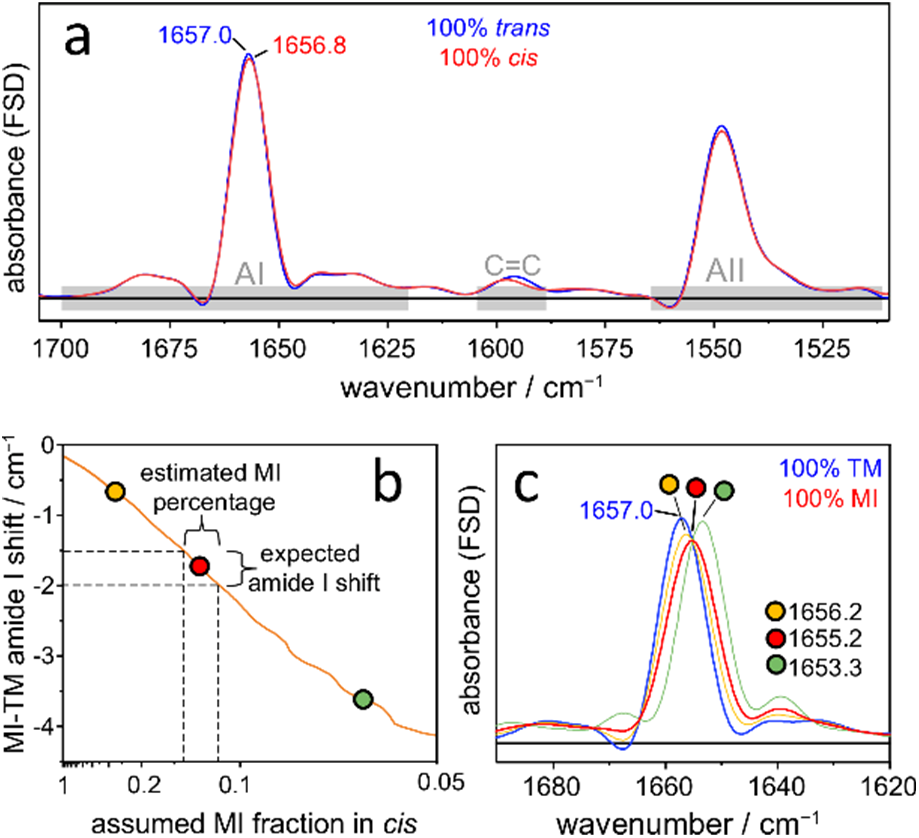
Mixture of TM and MI states in *cis* KCALP-azo. (a) FSD absorption spectrum of 100% *trans* KCALP-azo, adopting a 100% TM state (blue, reproduced from Fig. S4a). Estimated absorption spectrum of 100% *cis* KCALP-azo (red). The small downshift of the amide I, 0.2 cm^−1^, indicates that *cis* KCALP-azo exists in a TM and MI mixture. (b) Downshift of the amide I maximum between the 100% MI spectrum (estimated with Eq. 3) and the 100% TM spectrum (taken from 100% *trans* KCALP-azo) as a function of the value assumed for 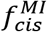 (fraction of *cis* KCALP-azo in the MI state). From the expected MI-TM amide I downshift, 1.5-2.0 cm^−1^ (dashed horizontal lines), we obtained an estimate of 0.14-0.11 for 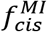 (dashed vertical lines). Three values for 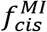 are labelled: optimal (red), overestimated (orange) and underestimated (green). (c) Amide I spectrum of KCALP-azo in a 100% TM state (blue, same as in (a)) and in a 100% MI state, the latter reconstructed assuming *cis* KCALP-azo to be 88% TM and 12% MI. Reconstructions of the 100% MI spectrum assuming 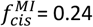 (orange) or 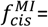 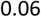 (green), display an amide I downshift smaller (0.8 cm^−1^) or larger (3.7 cm^−1^) than expected.

However, the amide I band downshifts by only 0.16 cm^−1^. In contrast, for LAH_4_ the amide I downshifts by ~1.9 cm^−1^ between the MI and TM states (Fig. S10), and our band decomposition analysis for KCALP-azo suggests that its amide I downshifts by ~1.6 cm^−1^ between the MI and TM states (Fig. 4e). This discrepancy indicates that the absorption spectrum of *cis* KCALP-azo arises from a mixture of TM and MI states:

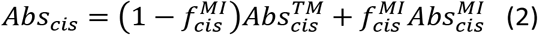

To estimate the amide I spectrum of *cis* KCALP-azo in the MI state, 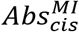, we assumed that the amide I spectrum depends on the membrane topology adopted by the peptide but it is not significantly affected by the isomeric state adopted by azobenzene, *i.e.*, 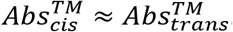. After considering that *trans* KCALP-azo is 100% TM, *i.e.*, 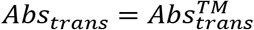, and rearranging Eq. 2, we arrived to:

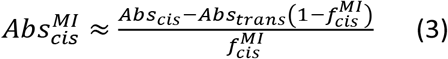

Using this equation we estimated 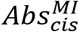 for different values of 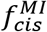, the fraction of *cis* KCALP-azo in a MI state. Figure 6b plots how much the amide I downshifts between the 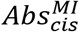 and Abs_*trans*_ as a function of the value assumed for 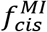. Because we expect this downshift to be around 1.5-2.0 cm^−1^, we deduced that 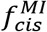 takes a value between 0.11 and 0.14 (see Fig. 6b, dashed lines), *i.e*., ~11-14% of *cis* KCALP-azo adopts a MI state. As an illustration, Fig. 6c shows the reconstructed amide I spectrum for the MI state, estimated using 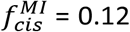 (red trace). The same plot shows, for reference, a reconstruction using a twice-larger (orange trace) or a twice-smaller (green trace) value.

We obtained an additional estimation for 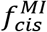. The area of the negative band at 1657.2 cm^−1^ in the light-induced FTIR difference spectrum (Fig. 4e and Table S5) is 60 times smaller than the area of the positive band at 1657.0 cm^−1^ in the FTIR absorption spectrum of *trans* KCALP-azo (Fig. 3c and Table S4). From this result, and taking into account that only 20% of the azobenzene were photoisomerized, we inferred that just 8% of *cis* KCALP-azo adopts a MI topology. Thus, two different lines of reasoning lead us to the conclusion that *cis* KCALP-azo exists as a mixture of ~90% TM and ~10% MI states.

## General discussion

The coupling of azobenzene to Cys side chains has been one of the approaches used in the past to introduce structural perturbations on soluble helical polypeptides with light.^14,20–22^ Here, we have extended this approach to KCALP-azo, a 24 residues-long TM-helical peptide. First, we characterized the structure and orientation of KCALP-azo in its dark-adapted state (~100% *trans* isomer). We have confirmed that KCALP-azo is indeed a TM and highly helical (~85%) peptide (Fig. 3). In addition, we found that *trans* KCALP-azo displays two types of helical segments with distinct properties. The dominant helix type (~70%) displays an amide I frequency at 1657 cm^−1^, characteristic for α-helices in TM proteins and peptides,^35,45^ and shows an average tilt of 36 ± 4° (Fig. 3, and ① in Fig. 7b). We assign it to the core of the TM helix of KCALP-azo (① in Fig. 7a). The minor helix type (~15%) displays an amide I frequency at 1646 cm^−1^, characteristic for helices forming bifurcated H-bonds with water molecules,^46,47^ and shows an average tilt of 49 ± 3° (Fig. 3, and ② in Fig. 7b). Thus, it likely corresponds to short segments at the two ends of the TM helix (② in Fig. 7a).

**Figure. 7.**
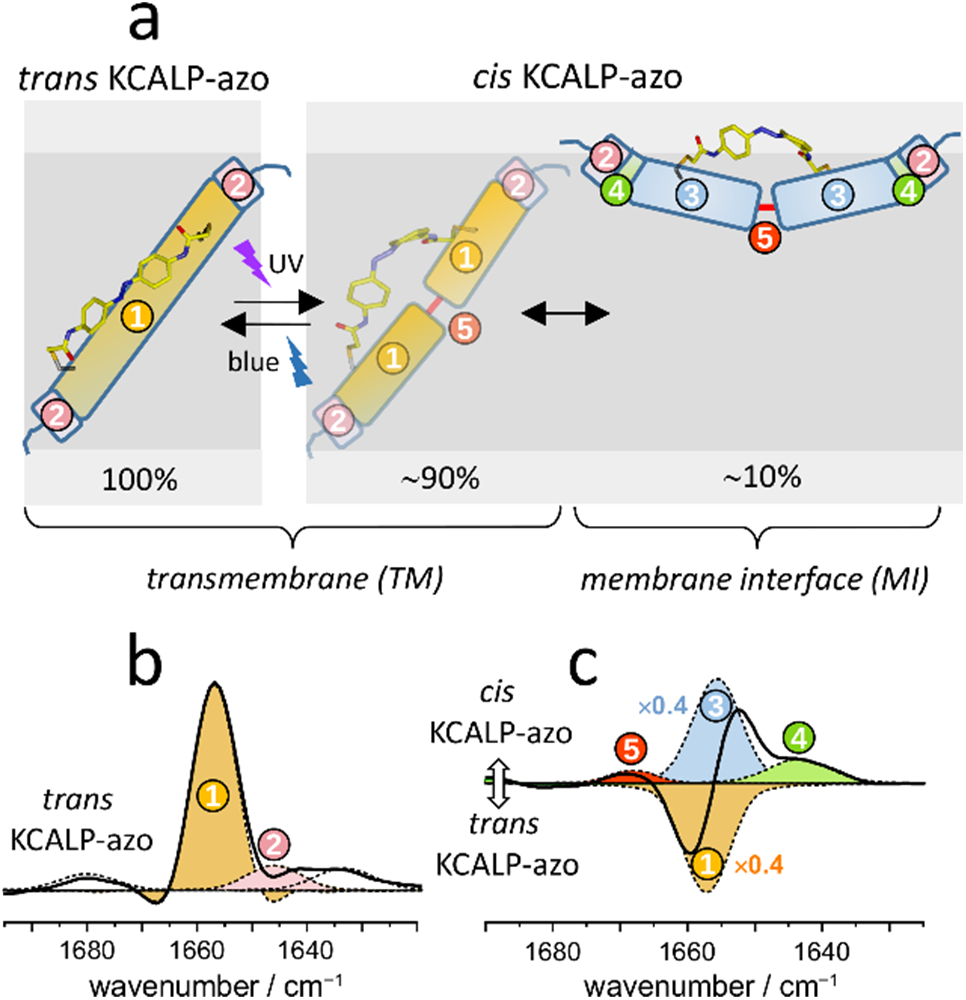
Structural models of the photoswitchable peptide KCALP-azo in *trans*-azobenzene-TM and *cis*-azobenzene-TM/MI states. (a) Structure and membrane topology of KCALP-azo when the azobenzene group is in either *trans* (left) or *cis* (right) isomeric state. The helical peptide structures are illustrated as cylinders, tilted according to the experimental results (azimuthal rotation angles are tentative). (b) Band-narrowed FTIR absorption spectrum of *trans* KCALP-azo, with its component bands resolved by band-fitting. (c) Band-narrowed light-induced FTIR difference spectrum of KCALP-azo (*cis* minus *trans*), with its component bands resolved by band-fitting. Related structural/spectral elements in (a-c) are labelled using common colours and number codes.

We next studied changes in the structure and orientation of KCALP-azo after being exposed to alternating UV and blue light pulses, which converts 20% of the azobenzene groups from a *trans* to a *cis* conformation. In the FTIR difference spectrum, the band at 1657.2 cm^−1^ represents 99% of the negative amide I area (① in Fig. 7c, see also Fig. 4e and Table S5), and shows a frequency and width very similar to the amide I band at 1657.0 cm^−1^ in the absorbance spectrum of KCALP-azo (① in Fig. 7b, see also Table S4). This high similitude indicates that photoisomerization of azobenzene affects almost exclusively to the hydrophobic core-segment of the TM helix of KCALP-azo. On the other hand, we observed three positive amide I bands in the light-induced FTIR difference spectrum of KCALP-azo (Fig. 4e). The main positive band (1655.6 cm^−1^) accounts for 86% of the amide I area (③ in Fig. 7c, see also Fig. 4e and Table S5), and it originates from the amide I vibration of a helix tilted by ~79 ± 8° with respect the membrane normal (Fig. 5). Thus, as a result of azobenzene photoisomerization, the central TM hydrophobic segment of KCALP-azo changes, both in amide I frequency (from ~1657.2 cm^−1^ to ~1655.6 cm^−1^) and in average tilt (from 36 ± 4° to 79 ± 8°), as shown in Fig. 7 (see ① to ③).

Although the helix tilt angle of a peptide does not directly define its topology in the membrane, it certainly constrains it. Considering the hydrophobic thickness of POPC in fluid phase, ~29 Å,^69^ a TM topology with an average helix tilt of 79 ± 8° can be discarded just from energetic considerations, as it would involve placing at least two of the four positively charged Lys residues of KCALP-azo (Fig. 1b) within the hydrophobic region of the membrane. The helix tilt of ~79 ± 8° for the MI state of *cis* KCALP-azo agrees well with that of mastoparan X, a helical peptide adopting MI topology, with a tilt angle determined to be 80 ± 5° by NMR^64^ and 76 ± 7° by FTIR spectroscopy.^65^ In addition, the shift in the amide I frequency between bands ① and ③ (Fig. 7c), is very similar to the shift that we observed between the TM and MI states of the pH-sensitive helical peptide LAH_4_ (Fig. S10).

The migration of KCALP-azo to the interface of the membrane goes accompanied by an enlargement of its hydrated helical segment (see ④, Fig. 7a), as indicated by the positive band at 1643.5 cm^−1^ (④ in Fig. 7c, see also Table S5). Finally, we also have a small positive band at 1668 cm^−1^ (⑤ in Fig. 7c, see also Table S5). Although the assignment of this band is tentative, given its wavenumber it might originate from weakly H-bonded backbone peptide groups formed as result of the distortion of the TM helix by *trans*-to-*cis* photoisomerization. We postulate that such distortion might localize near the centre of the helix core (⑤ in Fig. 7a), allowing for a bend in the TM helix in response to *trans*-to-*cis* isomerization of azobenzene.

We lack experimental information about the azimuthal rotation angle displayed by the TM and MI helices of KCALP-azo. While this angle might not be very relevant for the TM state, for the MI state it will define which residues face the interface region of the membrane, as well as the location of the azobenzene group. Direct information about the actual azimuthal rotation angle of KCALP-azo could be obtained in the future by polarized FTIR experiments in combination with a suitable site specific labelling strategy, following the work by Arkin and coworkers.^70^ At present, it is most reasonable to assume that the azobenzene group faces the membrane interface, as illustrated in Fig. 7.

The changes reported here for KCALP-azo are drastically different from those reported before for photoswitchable soluble helical peptides. As an example, azobenzene *trans*-to-*cis* photoisomerization was shown to almost completely unfold a 16 residue-long soluble helical peptide, reducing its helix content from 93% (*trans*) to 34% (*cis*).^19^ Why does a photoswitchable hydrophobic helical peptide reconstituted in membranes respond so differently to light? We have to take into account that the energetic cost for breaking a backbone amide H-bond in the hydrophobic core of lipidic membranes, +4-5 kcal/mol,^4,71^ is much higher than in aqueous solution, +0.5-1 kcal/mol.^71^ This makes unlikely that photoisomerization of azobenzene could easily induce the unfolding of a TM helix to any degree close to that reported before for soluble peptides, and agrees with our observation that KCALP-azo largely preserves its helical structure upon *trans*-to-*cis* photoisomerization.

In order to understand why azobenzene photoisomerization induced a change in the membrane topology of KCALP-azo, we need to be aware that the energetically closest state to a TM helix is not an unfolded TM state but a helix located at the interface of the membrane.^4^ From the sequence of KCALP, and using the hydrophobicity scales from Wimley and White^72,73^ implemented in the MPEx tool,^74^ we estimated the free energy difference between its TM and an MI states, hereafter ΔGoTM-MI, to be −4.5 kcal/mol. In this estimation we took into account only residues from Leu4 to Leu21, because we expect the three residues at either peptide end (GKK at the N-terminus and KKA at the C-terminus) to face the interface of the membrane regardless of the membrane topology adopted by KCALP. For KCALP-azo, we could not estimate ΔG°_TM-MI_ from MPEx, but it is plausible for KCALP and *trans* KCALP-azo to share a similar ΔG°_TM-MI_ when considering that both peptides show a similar elution time (*i.e.*, hydrophobicity) on an HPLC C8 column (see Fig. S11a and Fig. S13a). The ΔG°_TM-MI_ value provided by MPEx is sufficiently favourable to ensure that virtually 100% of KCALP (and *trans* KCALP-azo) adopts a TM topology, yet sufficiently low for changes in noncovalent interactions to significantly modulate the TM propensity of this peptide. The experimentally estimated TM/MI equilibrium constant for *cis* KCALP-azo, ~90/10 = 9, indicates a ΔG°_TM-MI_ of −1.3 kcal/mol. This value suggest that azobenzene *trans*-to-*cis* photoisomerization is able to increase ΔG°_TM-MI_ of KCALP-azo by ~3.2 kcal/mol (from −4.5 to −1.3 kcal/mol). Because breaking an intramolecular H-bond costs 3-4 kcal/mol less at the membrane interface than at the hydrophobic region of the membrane,^4^ we propose that azobenzene photoisomerization breaks (or weakens) one or more intramolecular H-bonds of KCALP-azo, favouring the peptide migration to the MI. In addition, we should not forget that the higher polarity of *cis* azobenzene compared with the *trans* isomer^21^ might also contribute to stabilize the MI state. Indeed, *cis* KCALP-azo is more polar than *trans* KCALP-azo, as indicated by the earlier elution time of the former in an HPLC C8 column (Fig. S7b).

Although our experimental data clearly points to a change in the membrane topology of KCALP-azo upon azobenzene photoisomerization, this change in topology is apparently far from complete. We roughly estimated that only ~10% of those peptides with the azobenzene group in *cis* conformation experienced a change in membrane topology (Fig. 6). Consequently, the measured static light-induced FTIR difference spectrum of KCALP-azo (Fig. 4c) actually represents a mixture of two different spectra:(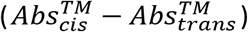 and 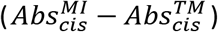). Future time-resolved experiments should be able to disentangle these two spectral contributions, because structural changes that immediately follow the photoisomerization of the azobenzene 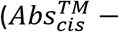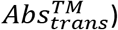 are expected to occur much earlier than the changes associated with variations of the membrane topology 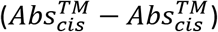.

Under our experimental illumination conditions only 20% of KCALP-azo photoisomerized from the *trans* to the *cis* state. From this 20%, only ~10% changed membrane topology. Thus, only ~2% of the total peptide switched with light from a TM to a MI state in our experiments. This small percentage did not represent a serious limitation for our sensitive FTIR difference spectroscopic experiments. In fact, future time-resolved FTIR studies aiming at revealing the mechanism of membrane insertion of helical peptides might benefit from a small population of “perturbed” peptides, limiting unwanted changes in the properties of the lipidic membrane. Indeed, in our static light-induced FTIR difference spectra we could observe only tiny bands assignable to lipid vibrations (Fig. 4c), indicating that the main physical properties of the POPC bilayers, like phase and fluidity, were not significantly altered by the change of topology of ~2% of KCALP-azo. However, for studies relying on other spectroscopic techniques, such as solid-state NMR or oriented CD, a much larger perturbation in the TM/MI equilibrium is desirable. This applies as well for future molecular designs aiming to exploit for practical purposes the optical control of the membrane topology of peptides with photoswitches. Possible strategies to enhance the formation of the MI state with light might include changes in the chemical nature or coupling of the photoswitch, but also an increase in the polarity of peptide. As an example, exchanging two Leu residues by Ala in the primary structure of KCALP should increase ΔG°_TM-MI_ by ~2 kcal/mol.^74^ Extrapolating this prediction to KCALP-azo leads to ΔG° of −2.5 kcal/mol for the *trans* conformation (TM/MI of ~99%/1%) and +0.7 kcal/mol for the *cis* conformation (TM/MI ~25%/75%), *i.e.*, a potential 74% change of the membrane topology with light. We are currently exploring this strategy, with positive results.

## Conclusions

In the present work, we have expanded the application of photoswitches in a new direction: the reversible optical control of the membrane topology of helical peptides. The possibility of repetitively perturb with light the membrane interface / transmembrane equilibrium of a peptide will make possible currently unfeasible experiments. One example are time-resolved infrared experiments, rich in structural and dynamic information, but which require laser pulses as a trigger (for the highest temporal resolution) as well as extensive data averaging (tremendously facilitated when the system is reversible).^75,76^ Thus, the here developed photoswitchable peptide will be a key step forward in the road of understanding the sequence of events that allow peptides (and protein fragments) to spontaneously insert into or move out of membranes.

The optical control of the membrane topology of peptides might be also the starting point for interesting applications. For instance, the pH-sensitive peptide pHLIP can deliver polar molecules towards the interior of cells in response to a membrane interface-to-transmembrane reorientation change induced at low pH, a property used to target drugs to tumour cells.^77^ Likewise, light-control of the membrane topology of peptides could allow for the delivery of small molecules by the same mechanism as for pHLIP, *i.e.*, transport of small cargo molecules without peptide penetration into the cell, although with the added benefits in temporal and spatial resolution that light provides.

## Experimental Section

### Peptide synthesis and purification

KCALP (Fig. 1) and LAH_4_ (KKALLALALHHLAHLALHLALALKKA-NH_2_)^78^ were prepared by a stepwise micro-wave assisted solid-phase peptide synthesis on a Liberty Blue synthesizer using the standard Fmoc/tBu strategy. Both peptides were constructed on a Rink Amide MBHA resin using a 5-fold molar excess of Fmoc-protected amino acids for chain elongation, acetic anhydride/pyridine 1/2 (v/v) for N-terminus acetylation, and trifluoroacetic acid (TFA)/thioanisole/1,2-ethanedithiol/anisole 90/5/3/2 (v/v/v/v) for final cleavage from the resin and removal of side chain protecting groups. Crude peptides were purified by semi-preparative RP-HPLC on a Phenomenex Luna C8(2) column (10 µm, 250 mm x 21.2 mm) using an acetonitrile/water gradient in 0.1% TFA. After lyophilization, the TFA counterions, which hamper physicochemical characterization by FTIR spectroscopy, were exchanged to chloride using a strong base anion exchange resin (AMBERLITE™ IRA402 Cl). High-resolution electrospray-ionization mass spectra (HRMS-ESI+) confirmed the identity of the peptides. KCALP: m/z for C_115_H_214_N_29_O_25_S_2_ [M+3H]^3+^ calculated 821.8597, found 822.1969 (Fig. S11c); LAH_4_: m/z for C_132_H_233_N_39_O_26_ [M+4H]^4+^ calculated 695.2022, found 695.4522 (Fig. S12c). HRMS-ESI+ was recorded in positive ion mode on a micrOTOF-Q spectrometer (Bruker), with sodium formate as an external reference. The purity of the peptides (>95%) was checked by analytical HPLC on a Phenomenex Luna C8(2) column (5 μm, 250 mm × 4.6 mm), as explained in Fig. S11b (for KCALP) and Fig. S12b (for LAH_4_).

### Synthesis of KCALP-azo, BCA and ThioAzo

KCALP (16 mg, 6.5 mmol) and the reducing agent tris(carboxyethyl)phosphine (7.45 mg, 26 mmol) were mixed and incubated overnight under an inert atmosphere of argon in a mixture of trifluoroethanol (TFE)/100 mM ammonium bicarbonate buffer (1/1 v/v, 10.2 mL, pH 8.5). A solution of BCA cross-linker (9.46 mg, 26 mmol) in dimethyl sulfoxide (2.6 mL) was added, and the mixture was stirred at 40 °C in the dark until completion of the reaction (monitored by MALDI-TOF mass spectrometry). The cross-linked peptide was purified by RP-HPLC and characterized by HRMS-ESI+: m/z for C_131_H_226_N_33_O_27_S_2_ [M+3H]^3+^ calculated 919.2250, found 919.5601 (Fig. S13c). The TFA counterions were exchanged for chloride as described for KCALP, and the purity of the product (>95%) was checked by analytical HPLC on a Phenomenex Luna C8(2), as shown in Fig. S13b. BCA (4,4’-bis(chloroacetamide)azobenzene) was synthesized according to literature, with spectral properties matching previously reported values.^32^ ThioAzo ((E)-N,N’-(diazene-1,2-diylbis(4,1-phenylene))bis(2-((2-hydroxyethyl)thio)acetamide) was synthesized as described above for KCALP-azo, except that 2-mercaptoethanol was used for the reaction with BCA instead of KCALP. The pure product was characterized by HRMS-ESI+ (Fig. S16) as well as by ^1^H and ^13^C NMR (Fig. S14 and S15), the latter recorded on a Bruker AVANCE III HD spectrometer (300 MHz).

### Peptide reconstitution in POPC membranes

Lyophilized peptides (KCALP, KCALP-azo and LAH_4_) were weighted, dissolved in TFE and mixed with in the appropriate amount of the POPC lipid (Avanti) in chloroform/methanol 2/1 (v/v), to a target lipid/peptide ratio of 20 (mol/mol), and dried in N2 atmosphere (followed by 2 hours of vacuum). Then, buffer was added, and the mixture was vigorously vortexed to form large multilamellar vesicles (LMV), washed 3 times by ultracentrifugation (50,000 RPM for 50 minutes), and finally resuspended in the appropriate volume of buffer (2 mM sodium phosphate at pH 7). Likewise, we prepared LMVs containing the ThioAzo compound alone, as well as LMVs containing equimolar amounts of KCALP and ThioAzo.

### Preparation and characterization of oriented films

We dried KCALP or KCALP-azo reconstituted in POPC vesicles (10-15 μL) at ambient humidity over BaF_2_ windows, forming films of ~6-8 mm diameter. For ATR experiments, ~2-4 μL of KCALP-azo and LAH_4_ reconstituted in POPC vesicles were dried on a silicon with 3 total reflections (DuraDisk, Czitek). These films were hydrated by the atmosphere created by drops of water/glycerol (see Fig. 3a, left).^79^ We estimated the final number of water molecules per peptide as described in Fig. S3. This ratio was ~950 in hydrated films used for unpolarized light-induced FTIR experiments (Fig. 4) and for polarized light-induced FTIR experiments by ATR (Fig. 5a), and ~450 for the rest of measurements (Fig. 3 and Fig. 5b). Both hydration levels provided similar results (see Fig. S4).

### FTIR experiments

Unpolarized FTIR absorption spectra were collected at 2 cm^−1^ spectral resolution, and light-induced unpolarized spectra at 4 cm^−1^ resolution, using either a Vertex 80v or a Vertex 80 FTIR spectrometer (Bruker) equipped with a photovoltaic MCT detector. Polarized absorption spectra were collected at 4 cm^−1^ resolution on a Nicolet 5700 (ThermoFisher) FTIR spectrometer equipped with a photoconductive MCT, using a BaF_2_ holographic wire grid polarizer (Thorlabs) mounted on a motorized rotational stage (Thorlabs, PRM1/MZ8). Polarized light-induced FTIR difference spectra were measured similarly, but on a Vertex 80 FTIR spectrometer. Illumination during transmission experiments was achieved using @365 nm and @447 nm LEDs, with power densities at the sample of ~400 mW/cm^2^ and ~200 mW/cm^2^, as measured with a powermeter (Thorlabs, PM160T). For ATR experiments the samples were illuminated from the top, with @365 nm and @455 nm LEDS coupled to optical fibers and power densities at the sample of ~100 mW/cm^2^ and ~175 mW/cm^2^. All the experiments were conducted at room temperature (~25 °C).

### UV-Vis experiments

UV-Vis absorption spectra of the pure *cis* and *trans* isomers of KCALP-azo were recorded by a Waters 2998 photodiode array detector built into the analytical HPLC equipment (Fig S7c). This allowed to determine the percentage of each isomer in solution (1/1 (v/v) acetonitrile/water +0.1% TFA), as described in Fig. S7. UV-Vis absorption spectra of hydrated films of KCALP-azo in POPC were measured in a home-made optical setup using an array detector (Flame-S-UV-Vis, OceanOptics). The absorption spectra of KCALP-azo were corrected from scattering using an absorption spectrum of a KCALP film as a blank.

### Spectral analysis

Fourier self-deconvolution (FSD) and band-fitting of FSD spectra were performed using homemade scripts running in MATLAB.^39,40^ For FSD we used a narrowing factor (*k*) of 2.0 and a Lorentzian width (γ_L_) of 18 cm^−1^, except in Fig. 4d,e (*k* = 2.0 and γ_L_ = 12 cm^−1^) and in the amide A range in Fig. 3d (*k* = 2.0 and γ_L_ = 50 cm^−1^). We fitted FSD spectra using FSD-modified Voigtian bands, using a method that takes into account bandshape changes introduced by FSD.^39,40^

### Determination of helical tilts

We applied Eq.1 to estimate *S*_*β*_, using distributed values for α, *n*_2_, *S*_*ms*_, *S*_*φ*_ and *R* that accounted for their uncertainty. We obtained probability distributions for *S*_*β*_ by generating 10^6^ random samples, binning the result and normalizing the area to 1. For the tilt angle of the sample window, α, we considered a uniform distribution from 48° to 52°. For *n*_2_ we used a uniform distribution from 1.45 to 1.55 for dry films, and from 1.35 to 1.45 for hydrated films. Regarding *S*_*φ*_, we assumed values of φ to follow a uniform distribution from 27° and 34° for the amide A vibration, from 38° and 41.5° for the amide I, and from 68° and 76° for the amide II.^41,62,63^ The value of *S*_*ms*_ in hydrated films was estimated to be 0.88 ± 0.03 by comparing the order parameter for the *ν*_*s*_CH vibration determined by polarized FTIR experiments, −0.145 ± 0.005, with the one deduced using a published correlation between its order parameter and its *ν*_*s*_CH frequency:80 −0.170 ± 0.005 for a 2853.45 ± 0.05 cm^−1^ *ν*_*s*_CH frequency. We assumed that the same value for *S*_*ms*_ applies to the dry films. Finally, we generously assumed the experimental value of *R* to have an error of ± 0.02. To estimate the average helical tilt, ⟨*β*⟩, from *S*_*β*_ we used the approximation *S*_*β*_ ≈ (3cos^2^⟨*β*⟩ − 1)⁄2.

## Author Contributions

D.S. and V.A.L.-F. designed the research, M.G.-S., E.S.-A., L.S. and V.A.L.-F. performed experiments and analysed data, M.G.-S., E.S.-A., D.S., J.S. and V.A.L.-F. discussed the results and wrote the paper.

## Acknowledgements

We thank the financial support provided by the Spanish Ministry of Science and Innovation (MICINN) through projects BFU2016-768050-P, BFU2017-91559-EXP, PID2019-106103GB-I00 and CTQ2017-87372-P, and by the Valencian Government through the project PROMETEU/2019/066. V.A.L.-F acknowledges a Ramon y Cajal fellowship (RYC-2013-13114), M.G.-S. a predoctoral fellowship (BES-2017-080385) from the MICINN, and E.S.-A. a predoctoral fellowship from Universidad de La Rioja. V.A.L.-F. is in debt with Joachim Heberle for access to a Vertex 80v FTIR spectrometer in the early stage of this work, and thanks Mattia Saita and Franziska Sellnau for assistance during those measurements.

## References

(1) Park, E.; Rapoport, T. A. Mechanisms of Sec61/SecY-Mediated Protein Translocation Across Membranes. Annu. Rev. Biophys. 2012, 41 (1), 21–40. https://doi.org/10.1146/annurev-biophys-050511-102312.

(2) Brambillasca, S.; Yabal, M.; Makarow, M.; Borgese, N. Unassisted Translocation of Large Polypeptide Domains across Phospholipid Bilayers. J. Cell Biol. 2006, 175 (5), 767–777. https://doi.org/10.1083/jcb.200608101.

(3) Guha, S.; Ghimire, J.; Wu, E.; Wimley, W. C. Mechanistic Landscape of Membrane-Permeabilizing Peptides. Chem. Rev. 2019, 119 (9), 6040–6085. https://doi.org/10.1021/acs.chemrev.8b00520.

(4) White, S. H.; Wimley, W. C. Membrane Protein Folding and Stability: Physical Principles. Annu. Rev. Biophys. Biomol. Struct. 1999, 28, 319–365. https://doi.org/10.1146/annurev.biophys.28.1.319.

(5) Ulmschneider, J. P.; Ulmschneider, M. B. Molecular Dynamics Simulations Are Redefining Our View of Peptides Interacting with Biological Membranes. Acc. Chem. Res. 2018, 51 (5), 1106–1116. https://doi.org/10.1021/acs.accounts.7b00613.

(6) Hunt, J. F.; Rath, P.; Rothschild, K. J.; Engelman, D. M. Spontaneous, PH-Dependent Membrane Insertion of a Transbilayer Alpha-Helix. Biochemistry 1997, 36 (49), 15177–15192. https://doi.org/10.1021/bi970147b.

(7) Tang, J.; Gai, F. Dissecting the Membrane Binding and Insertion Kinetics of a PHLIP Peptide. Biochemistry 2008, 47 (32), 8250–8252. https://doi.org/10.1021/bi801103x.

(8) Andreev, O. A.; Karabadzhak, A. G.; Weerakkody, D.; Andreev, G. O.; Engelman, D. M.; Reshetnyak, Y. K. PH (Low) Insertion Peptide (PHLIP) Inserts across a Lipid Bilayer as a Helix and Exits by a Different Path. Proc. Natl. Acad. Sci. U. S. A. 2010, 107 (9), 4081–4086. https://doi.org/10.1073/pnas.0914330107.

(9) Schuler, E. E.; Nagarajan, S.; Dyer, R. B. Submillisecond Dynamics of Mastoparan X Insertion into Lipid Membranes. J. Phys. Chem. Lett. 2016, 7 (17), 3365–3370. https://doi.org/10.1021/acs.jpclett.6b01512.

(10) Roder, H.; Maki, K.; Cheng, H. Early Events in Protein Folding Explored by Rapid Mixing Methods. Chem. Rev. 2006, 106 (5), 1836–1861. https://doi.org/10.1021/cr040430y.

(11) Kubelka, J. Time-Resolved Methods in Biophysics. 9. Laser Temperature-Jump Methods for Investigating Biomolecular Dynamics. Photochem. Photobiol. Sci. 2009, 8 (4), 499–512. https://doi.org/10.1039/b819929a.

(12) Blanco-Lomas, M.; Samanta, S.; Campos, P. J.; Woolley, G. A.; Sampedro, D. Reversible Photocontrol of Peptide Conformation with a Rhodopsin-like Photoswitch. J. Am. Chem. Soc. 2012, 134 (16), 6960–6963. https://doi.org/10.1021/ja301868p.

(13) Kumita, J. R.; Smart, O. S.; Woolley, G. a. Photo-Control of Helix Content in a Short Peptide. Proc. Natl. Acad. Sci. 2000, 97 (8), 3803–3808. https://doi.org/10.1073/pnas.97.8.3803.

(14) Hamm, P.; Helbing, J.; Bredenbeck, J. Two-Dimensional Infrared Spectroscopy of Photoswitchable Peptides. Annu. Rev. Phys. Chem. 2008, 59, 291–317. https://doi.org/10.1146/annurev.physchem.59.032607.093757.

(15) Flint, D. G.; Kumita, J. R.; Smart, O. S.; Woolley, G. A. Using an Azobenzene Cross-Linker to Either Increase or Decrease Peptide Helix Content upon Trans-to-Cis Photoisomerization. Chem. Biol. 2002, 9 (3), 391–397. https://doi.org/10.1016/S1074-5521(02)00109-6.

(16) Woolley, G. A. Photocontrolling Peptide Alpha Helices. Acc. Chem. Res. 2005, 38 (6), 486–493. https://doi.org/10.1021/ar040091v.

(17) Garcia-Iriepa, C.; Gueye, M.; Leonard, J.; Martinez-Lopez, D.; Campos, P. J.; Frutos, L. M.; Sampedro, D.; Marazzi, M. A Biomimetic Molecular Switch at Work: Coupling Photoisomerization Dynamics to Peptide Structural Rearrangement. Phys. Chem. Chem. Phys. 2016, 18 (9), 6742–6753. https://doi.org/10.1039/C5CP07599H.

(18) Ihalainen, J. A.; Bredenbeck, J.; Pfister, R.; Helbing, J.; Chi, L.; van Stokkum, I. H. M.; Woolley, G. A.; Hamm, P. Folding and Unfolding of a Photoswitchable Peptide from Picoseconds to Microseconds. Proc. Natl. Acad. Sci. U. S. A. 2007, 104 (13), 5383–5388. https://doi.org/10.1073/pnas.0607748104.

(19) Bredenbeck, J.; Helbing, J.; Kumita, J. R.; Woolley, G. A.; Hamm, P. α-Helix Formation in a Photoswitchable Peptide Tracked from Picoseconds to Microseconds by Time-Resolved IR Spectroscopy. Proc. Natl. Acad. Sci. U. S. A. 2005, 102 (7), 2379–2384. https://doi.org/10.1073/pnas.0406948102.

(20) Albert, L.; Vázquez, O. Photoswitchable Peptides for Spatiotemporal Control of Biological Functions. Chem. Commun. 2019, 55 (69), 10192–10213. https://doi.org/10.1039/C9CC03346G.

(21) Beharry, A. A.; Woolley, G. A. Azobenzene Photoswitches for Biomolecules. Chem. Soc. Rev. 2011, 40 (8), 4422–4437. https://doi.org/10.1039/c1cs15023e.

(22) Szymański, W.; Beierle, J. M.; Kistemaker, H. A. V; Velema, W. A.; Feringa, B. L. Reversible Photocontrol of Biological Systems by the Incorporation of Molecular Photoswitches. Chem. Rev. 2013, 113 (8), 6114–6178. https://doi.org/10.1021/cr300179f.

(23) Hoersch, D.; Roh, S.-H.; Chiu, W.; Kortemme, T. Reprogramming an ATP-Driven Protein Machine into a Light-Gated Nanocage. Nat. Nanotechnol. 2013, 8 (12), 928–932. https://doi.org/10.1038/nnano.2013.242.

(24) Mutter, N. L.; Volarić, J.; Szymanski, W.; Feringa, B. L.; Maglia, G. Reversible Photocontrolled Nanopore Assembly. J. Am. Chem. Soc. 2019, 141 (36), 14356–14363. https://doi.org/10.1021/jacs.9b06998.

(25) Hoorens, M. W. H.; Szymanski, W. Reversible, Spatial and Temporal Control over Protein Activity Using Light. Trends Biochem. Sci. 2018, 43 (8), 567–575. https://doi.org/10.1016/J.TIBS.2018.05.004.

(26) Babii, O.; Afonin, S.; Ishchenko, A. Y.; Schober, T.; Negelia, A. O.; Tolstanova, G. M.; Garmanchuk, L. V.; Ostapchenko, L. I.; Komarov, I. V.; Ulrich, A. S. Structure–Activity Relationships of Photoswitchable Diarylethene-Based β-Hairpin Peptides as Membranolytic Antimicrobial and Anticancer Agents. J. Med. Chem. 2018, 61 (23), 10793–10813. https://doi.org/10.1021/acs.jmedchem.8b01428.

(27) Babii, O.; Afonin, S.; Berditsch, M.; Reiβer, S.; Mykhailiuk, P. K.; Kubyshkin, V. S.; Steinbrecher, T.; Ulrich, A. S.; Komarov, I. V. Controlling Biological Activity with Light: Diarylethene-Containing Cyclic Peptidomimetics. Angew. Chemie 2014, 126 (13), 3460–3463. https://doi.org/10.1002/ange.201310019.

(28) Yeoh, Y. Q.; Yu, J.; Polyak, S. W.; Horsley, J. R.; Abell, A. D. Photopharmacological Control of Cyclic Antimicrobial Peptides. ChemBioChem 2018, 19 (24), 2591–2597. https://doi.org/10.1002/cbic.201800618.

(29) Zhang, Y.; Lewis, R. N. A. H.; Henry, G. D.; Sykes, B. D.; Hodges, R. S.; McElhaney, R. N. Peptide Models of Helical Hydrophobic Transmembrane Segments of Membrane Proteins. 1. Studies of the Conformation, Intrabilayer Orientation, and Amide Hydrogen Exchangeability of Ac-K2-(LA)12-K2. Biochemistry 1995, 34 (7), 2348–2361. https://doi.org/10.1021/bi00007a031.

(30) Ulmschneider, M. B.; Doux, J. P. F.; Killian, J. A.; Smith, J. C.; Ulmschneider, J. P. Mechanism and Kinetics of Peptide Partitioning into Membranes from All-Atom Simulations of Thermostable Peptides. J. Am. Chem. Soc. 2010, 132 (10), 3452–3460. https://doi.org/10.1021/ja909347x.

(31) Strandberg, E.; Esteban-Martín, S.; Ulrich, A. S.; Salgado, J. Hydrophobic Mismatch of Mobile Transmembrane Helices: Merging Theory and Experiments. Biochim. Biophys. Acta - Biomembr. 2012, 1818 (5), 1242–1249. https://doi.org/10.1016/j.bbamem.2012.01.023.

(32) Pozhidaeva, N.; Cormier, M. E.; Chaudhari, A.; Woolley, G. A. Reversible Photocontrol of Peptide Helix Content: Adjusting Thermal Stability of the Cis State. Bioconjug. Chem. 2004, 15 (6), 1297–1303. https://doi.org/10.1021/bc049855h.

(33) Krimm, S.; Bandekar, J. Vibrational Spectroscopy and Conformation of Peptides, Polypeptides, and Proteins. In Advances in Protein Chemistry; Academic Press, 1986; Vol. 38, pp 181–364. https://doi.org/10.1016/S0065-3233(08)60528-8.

(34) Mantsch, H. H.; McElhaney, R. N. Phospholipid Phase Transitions in Model and Biological Membranes as Studied by Infrared Spectroscopy. Chem. Phys. Lipids 1991, 57 (2–3), 213–226. https://doi.org/10.1016/0009-3084(91)90077-O.

(35) Tamm, L. K.; Tatulian, S. A. Infrared Spectroscopy of Proteins and Peptides in Lipid Bilayers. Q. Rev. Biophys. 1997, 30 (4), 365–429. https://doi.org/10.1016/S0006-3495(95)80150-5.

(36) Venyaminov SYu; Kalnin, N. N. Quantitative IR Spectrophotometry of Peptide Compounds in Water (H_2_O) Solutions. II. Amide Absorption Bands of Polypeptides and Fibrous Proteins in α-, β-, and Random Coil Conformations. Biopolymers 1990, 30 (13–14), 1259–1271. https://doi.org/10.1002/bip.360301310.

(37) Kauppinen, J. K.; Moffatt, D. J.; Mantsch, H. H.; Cameron, D. G.; Spectroscopy, R. Fourier Self-Deconvolution: A Method for Resolving Intrinsically Overlapped Bands. Appl. Spectrosc. 1981, 35 (3), 271–276. https://doi.org/10.1366/0003702814732634.

(38) Pfister, R.; Ihalainen, J.; Hamm, P.; Kolano, C. Synthesis, Characterization and Applicability of Three Isotope Labeled Azobenzene Photoswitches. Org. Biomol. Chem. 2008, 6 (19), 3508. https://doi.org/10.1039/b804568b.

(39) Lórenz-Fonfría, V. A.; Padrós, E. Curve-Fitting of Fourier Manipulated Spectra Comprising Apodization, Smoothing, Derivation and Deconvolution. Spectrochim. Acta. A. Mol. Biomol. Spectrosc. 2004, 60 (12), 2703–2710. https://doi.org/10.1016/j.saa.2004.01.008.

(40) Lórenz-Fonfría, V. A.; Padrós, E. Curve-Fitting Overlapped Bands: Quantification and Improvement of Curve-Fitting Robustness in the Presence of Errors in the Model and in the Data. Analyst 2004, 129 (12), 1243–1250. https://doi.org/10.1039/b406581f.

(41) Marsh, D.; Müller, M.; Schmitt, F. J. Orientation of the Infrared Transition Moments for an Alpha-Helix. Biophys. J. 2000, 78 (5), 2499–2510. https://doi.org/10.1016/S0006-3495(00)76795-6.

(42) Rahmelow, K.; Hübner, W.; Ackermann, T. Infrared Absorbances of Protein Side Chains. Anal. Biochem. 1998, 257 (1), 1–11. https://doi.org/10.1006/abio.1997.2502.

(43) Kubelka, J.; Keiderling, T. A. Ab Initio Calculation of Amide Carbonyl Stretch Vibrational Frequencies in Solution with Modified Basis Sets. 1. N - Methyl Acetamide. J. Phys. Chem. A 2001, 105 (48), 10922–10928. https://doi.org/10.1021/jp013203y.

(44) Lorenz-Fonfria, V. A. Infrared Difference Spectroscopy of Proteins: From Bands to Bonds. Chem. Rev. 2020, 120 (7), 3466–3576. https://doi.org/10.1021/acs.chemrev.9b00449.

(45) Goormaghtigh, E.; Cabiaux, V.; Ruysschaert, J.-M. Determination of Soluble and Membrane Protein Structure by Fourier Transform Infrared Spectroscopy. III. Secondary Structures. Subcell. Biochem. 1994, 405–450. https://doi.org/10.1007/978-1-4615-1863-1_10.

(46) Walsh, S. T. R. R.; Cheng, R. P.; Wright, W. W.; Alonso, D. O. V; Daggett, V.; Jane, M.; Degrado, W. F.; Vanderkooi, J. M.; Grado, W. F. D. E. The Hydration of Amides in Helices; a Comprehensive Picture from Molecular Dynamics, IR, and NMR. Protein Sci. 2003, 12 (3), 520–531. https://doi.org/10.1110/ps.0223003.

(47) Lórenz-Fonfría, V. A.; Bamann, C.; Resler, T.; Schlesinger, R.; Bamberg, E.; Heberle, J. Temporal Evolution of Helix Hydration in a Light-Gated Ion Channel Correlates with Ion Conductance. Proc. Natl. Acad. Sci. U. S. A. 2015, 112 (43), E5796–804. https://doi.org/10.1073/pnas.1511462112.

(48) Ludlam, C. F.; Arkin, I. T.; Liu, X. M.; Rothman, M. S.; Rath, P.; Aimoto, S.; Smith, S. O.; Engelman, D. M.; Rothschild, K. J. Fourier Transform Infrared Spectroscopy and Site-Directed Isotope Labeling as a Probe of Local Secondary Structure in the Transmembrane Domain of Phospholamban. Biophys. J. 1996, 70 (4), 1728–1736. https://doi.org/10.1016/S0006-3495(96)79735-7.

(49) Dave, N.; Lórenz-Fonfría, V. A.; Leblanc, G.; Padrós, E. FTIR Spectroscopy of Secondary-Structure Reorientation of Melibiose Permease Modulated by Substrate Binding. Biophys. J. 2008, 94 (9), 3659–3670. https://doi.org/10.1529/biophysj.107.115550.

(50) Earnest, T. N.; Herzfeld, J.; Rothschild, K. J. Polarized Fourier Transform Infrared Spectroscopy of Bacteriorhodopsin. Transmembrane Alpha Helices Are Resistant to Hydrogen/Deuterium Exchange. Biophys. J. 1990, 58 (6), 1539–1546. https://doi.org/10.1016/S0006-3495(90)82498-X.

(51) DeLange, F.; Bovee-Geurts, P. H.; Pistorius, a M.; Rothschild, K. J.; DeGrip, W. J. Probing Intramolecular Orientations in Rhodopsin and Metarhodopsin II by Polarized Infrared Difference Spectroscopy. Biochemistry 1999, 38 (40), 13200–13209.

(52) Bechinger, B. Towards Membrane Protein Design: PH-Sensitive Topology of Histidine-Containing Polypeptides. J. Mol. Biol. 1996, 263 (5), 768–775. https://doi.org/10.1006/JMBI.1996.0614.

(53) Rothschild, K. J.; Clark, N. A. Polarized Infrared Spectroscopy of Oriented Purple Membrane. Biophys. J. 1979, 25 (3), 473–487. https://doi.org/10.1016/S0006-3495(79)85317-5.

(54) Nabedryk, E.; Breton, J. Orientation of Intrinsic Proteins in Photosynthetic Membranes. Polarized Infrared Spectroscopy of Chloroplasts and Chromatophores. Biochim. Biophys. Acta - Bioenerg. 1981, 635 (3), 515–524. https://doi.org/10.1016/0005-2728(81)90110-9.

(55) Esteban-Martín, S.; Giménez, D.; Fuertes, G.; Salgado, J. Orientational Landscapes of Peptides in Membranes: Prediction of 2H NMR Couplings in a Dynamic Context. Biochemistry 2009, 48 (48), 11441–11448. https://doi.org/10.1021/bi901017y.

(56) Ulmschneider, J. P.; Andersson, M.; Ulmschneider, M. B. Determining Peptide Partitioning Properties via Computer Simulation. J. Membr. Biol. 2011, 239 (1–2), 15–26. https://doi.org/10.1007/s00232-010-9324-8.

(57) Englander, S. W.; Mayne, L.; Kan, Z.-Y.; Hu, W. Protein Folding—How and Why: By Hydrogen Exchange, Fragment Separation, and Mass Spectrometry. Annu. Rev. Biophys. 2016, 45 (1), 135–152. https://doi.org/10.1146/annurev-biophys-062215-011121.

(58) Grdadolnik, J. Infrared Difference Spectroscopy: Part I. Interpretation of the Difference Spectrum. Vib. Spectrosc. 2003, 31 (2), 279–288. https://doi.org/10.1016/S0924-2031(03)00018-3.

(59) Bechinger, B.; Ruysschaert, J.-M.; Goormaghtigh, E. Membrane Helix Orientation from Linear Dichroism of Infrared Attenuated Total Reflection Spectra. Biophys. J. 1999, 76 (1), 552–563. https://doi.org/10.1016/S0006-3495(99)77223-1.

(60) Oh, K.-I.; Fiorin, G.; Gai, F. How Sensitive Is the Amide I Vibration of the Polypeptide Backbone to Electric Fields? ChemPhysChem 2015, 16 (17), 3595–3598. https://doi.org/10.1002/cphc.201500777.

(61) Lórenz-Fonfría, V. A.; Granell, M.; León, X.; Leblanc, G.; Padrós, E. In-Plane and Out-of-Plane Infrared Difference Spectroscopy Unravels Tilting of Helices and Structural Changes in a Membrane Protein upon Substrate Binding. J. Am. Chem. Soc. 2009, 131 (42), 15094–15095. https://doi.org/10.1021/ja906324z.

(62) Tsuboi, M. Infrared Dichroism and Molecular Conformation of α-Form Poly-γ-Benzyl-L-Glutamate. J. Polym. Sci. 1962, 59 (167), 139–153. https://doi.org/10.1002/pol.1962.1205916712.

(63) Marsh, D.; Páli, T. Infrared Dichroism from the X-Ray Structure of Bacteriorhodopsin. Biophys. J. 2001, 80 (1), 305–312. https://doi.org/10.1016/S0006-3495(01)76015-8.

(64) Whiles, J. A.; Brasseur, R.; Glover, K. J.; Melacini, G.; Komives, E. A.; Vold, R. R. Orientation and Effects of Mastoparan X on Phospholipid Bicelles. Biophys. J. 2001, 80 (1), 280–293. https://doi.org/10.1016/S0006-3495(01)76013-4.

(65) Tucker, M. J.; Getahun, Z.; Nanda, V.; DeGrado, W. F.; Gai, F. A New Method for Determining the Local Environment and Orientation of Individual Side Chains of Membrane-Binding Peptides. J. Am. Chem. Soc. 2004, 126 (16), 5078–5079. https://doi.org/10.1021/ja032015d.

(66) Mayo, D. J.; Sahu, I. D.; Lorigan, G. A. Assessing Topology and Surface Orientation of an Antimicrobial Peptide Magainin 2 Using Mechanically Aligned Bilayers and Electron Paramagnetic Resonance Spectroscopy. Chem. Phys. Lipids 2018, 213, 124–130. https://doi.org/10.1016/j.chemphyslip.2018.04.004.

(67) Strandberg, E.; Horn, D.; Reißer, S.; Zerweck, J.; Wadhwani, P.; Ulrich, A. S. 2H-NMR and MD Simulations Reveal Membrane-Bound Conformation of Magainin 2 and Its Synergy with PGLa. Biophys. J. 2016, 111 (10), 2149–2161. https://doi.org/10.1016/j.bpj.2016.10.012.

(68) Denschlag, R.; Schreier, W. J.; Rieff, B.; Schrader, T. E.; Koller, F. O.; Moroder, L.; Zinth, W.; Tavan, P. Relaxation Time Prediction for a Light Switchable Peptide by Molecular Dynamics. Phys. Chem. Chem. Phys. 2010, 12 (23), 6204–6218. https://doi.org/10.1039/b921803c.

(69) Kučerka, N.; Nieh, M. P.; Katsaras, J. Fluid Phase Lipid Areas and Bilayer Thicknesses of Commonly Used Phosphatidylcholines as a Function of Temperature. Biochim. Biophys. Acta - Biomembr. 2011, 1808 (11), 2761–2771. https://doi.org/10.1016/j.bbamem.2011.07.022.

(70) Arkin, I. T. Isotope-Edited IR Spectroscopy for the Study of Membrane Proteins. Curr. Opin. Chem. Biol. 2006, 10 (5), 394–401. https://doi.org/10.1016/j.cbpa.2006.08.013.

(71) Bolen, D. W.; Rose, G. D. Structure and Energetics of the Hydrogen-Bonded Backbone in Protein Folding. Annu. Rev. Biochem. 2008, 77 (1), 339–362. https://doi.org/10.1146/annurev.biochem.77.061306.131357.

(72) Wimley, W. C.; White, S. H. Experimentally Determined Hydrophobicity Scale for Proteins at Membrane Interfaces. Nat. Struct. Biol. 1996, 3 (10), 842–848. https://doi.org/10.1038/nsb1096-842.

(73) Wimley, W. C.; Creamer, T. P.; White, S. H. Solvation Energies of Amino Acid Side Chains and Backbone in a Family of Host-Guest Pentapeptides. Biochemistry 1996, 35 (16), 5109–5124. https://doi.org/10.1021/bi9600153.

(74) Snider, C.; Jayasinghe, S.; Hristova, K.; White, S. H. MPEx: A Tool for Exploring Membrane Proteins. Protein Sci. 2009, 18 (12), 2624–2628. https://doi.org/10.1002/pro.256.

(75) Lórenz-Fonfría, V. Infrared Difference Spectroscopy of Proteins: From Bands to Bonds. Chem. Rev. 120 (7), 3466–3576. https://doi.org/10.1021/acs.chemrev.9b00449.

(76) Kottke, T.; Lórenz-Fonfría, V. A.; Heberle, J. The Grateful Infrared: Sequential Protein Structural Changes Resolved by Infrared Difference Spectroscopy. J. Phys. Chem. B 2017, 121 (2), 335–350. https://doi.org/10.1021/acs.jpcb.6b09222.

(77) Wyatt, L. C.; Moshnikova, A.; Crawford, T.; Engelman, D. M.; Andreev, O. A.; Reshetnyak, Y. K. Peptides of PHLIP Family for Targeted Intracellular and Extracellular Delivery of Cargo Molecules to Tumors. Proc. Natl. Acad. Sci. U. S. A. 2018, 115 (12), E2811–E2818. https://doi.org/10.1073/pnas.1715350115.

(78) Vogt, T. C.; Bechinger, B. The Interactions of Histidine-Containing Amphipathic Helical Peptide Antibiotics with Lipid Bilayers. The Effects of Charges and PH. J. Biol. Chem. 1999, 274 (41), 29115–29121. https://doi.org/10.1074/jbc.274.41.29115.

(79) Noguchi, T.; Sugiura, M. Flash-Induced FTIR Difference Spectra of the Water Oxidizing Complex in Moderately Hydrated Photosystem II Core Films: Effect of Hydration Extent on S-State Transitions. Biochemistry 2002, 41 (7), 2322–2330.

(80) Kodati, V. R.; Lafleur, M. Comparison between Orientational and Conformational Orders in Fluid Lipid Bilayers. Biophys. J. 1993, 64 (1), 163–170. https://doi.org/10.1016/S0006-3495(93)81351-1.

