## Supplemental_Figures_Tables for "A photoswitchable helical peptide with light-controllable interface / transmembrane topology in lipidic membranes"

### Supplementary Information for
#### A photoswitchable hydrophobic helical peptide with light-controllable interface / transmembrane topology in lipidic membranes

Mónica Gutiérrez-Salazar, Eduardo Santamaría-Aranda, Louise Schaar, Jesús Salgado, Diego Sampedro, and Victor A. Lorenz-Fonfria

**This PDF file includes:**

Figures S1 to S16
Tables S1 to S5
SI References

**Supplementary Figures**

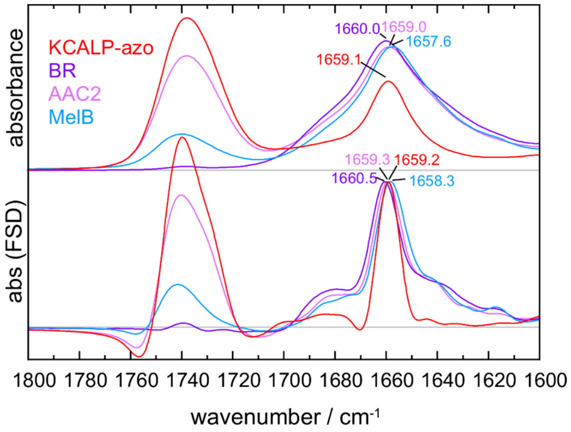


**Figure S1.** Normalized FTIR absorption spectrum of KCALP-azo reconstituted in POPC membranes compared to transmembrane proteins rich in α-helical structures: bacteriorhodopsin (BR) in purple membranes, yeast mitochondrial ATP/ADP carrier (AAC2) inhibited with atractyloside and reconstituted in egg PC/PA, and the bacterial $Na^+$-coupled melibiose permease (MelB) reconstituted in *E. coli* polar lipid extract, with helical contents of ~71% (pdb 1AT9),[1] ~77% (pdb 1OKC),[2] and ~62% (pdb 4M64),[3] respectively. The top panel display raw spectra, while bottom panel display band-narrowed spectra using Fourier self-deconvolution (FSD) $k = 2$ and $\gamma_L' = 18$ cm$^{-1}$. The wavenumber of the amide I peak maximum is indicated for each protein. Spectra in the top and the bottom panel are normalized to the same amide I peak intensity after FSD. Spectra of membrane proteins are reproduced from previous publications.[4–6] The percentage of helical structures have been obtained by applying the DSSP method[7] on structures deposited at the protein data bank, as analyzed by the web server 2Struc (https://2struccompare.cryst.bbk.ac.uk).[8]

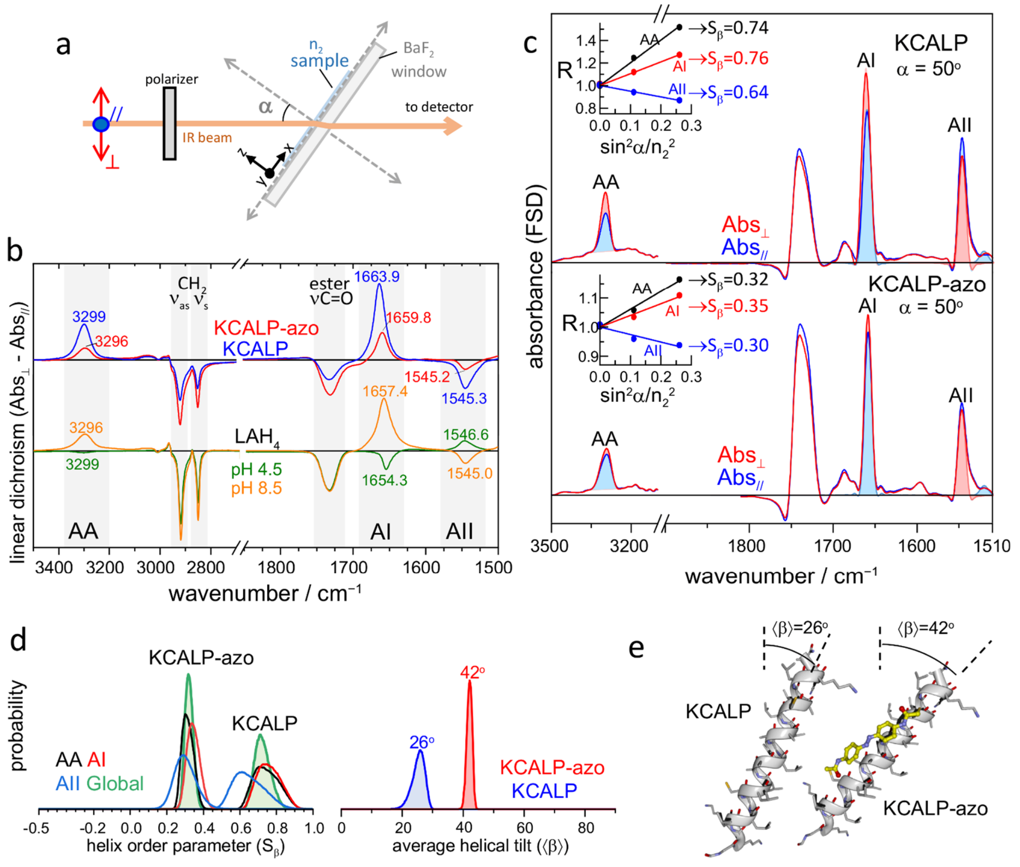


**Figure S2.** Membrane topology and helix tilt of KCALP and KCALP-azo in dried POPC membranes. (a) Sketch of the polarized FTIR setup by transmission, involving the use of polarized IR light and rotation of the sample window by an angle α. (b) Linear dichroism (LD) spectra of KCALP (blue trace) and KCALP-azo (red trace) for α = 50°, at 4 cm$^{-1}$ resolution (top). LD spectra for the pH-sensitive helical peptide $LAH_4$, which shows a membrane interface topology at pH 4.5 (green trace) and transmembrane topology at pH 8.5 (orange trace),[9] illustrating how the sign of the amide A, I and II bands in LD spectra depends on the peptide topology. Spectra for $LAH_4$ were measured by attenuated total reflection (ATR), and corrected for the dependence of the effective penetration on wavenumber.[10] (c) Band-narrowed polarized FTIR absorption spectra for α = 50°. FSD was performed with k = 2 and $\gamma_L$' = 50 cm$^{-1}$ above 3000 cm$^{-1}$, and below 2000 cm$^{-1}$ with k = 2 and $\gamma_L$' = 18 cm$^{-1}$. Band areas corresponding to helical structures, used to obtain dichroic ratios, are color-filled. For amide I and amide II vibrations these areas were measured by band-fitting, and for the amide A vibration by integration with a baseline. (Insets) Dichroic ratio (*R*) for amide A, I and II vibrations of helical structures in KCALP and KCALP-azo as a function of $sin^2\,\alpha/n_2^2$. (d, left) Probability distributions of the order parameter of the helical axis, $S_\beta$, of KCALP and KCALP-azo (see Experimental Section in the MS for more details). (d, right) Global probability for the average helical tilt, $\langle\beta\rangle$, of KCALP and KCALP-azo. (e) Schematic molecular model of the helical structure of KCALP and KCALP-azo, illustrating how the peptides tilt in dry POPC membranes.

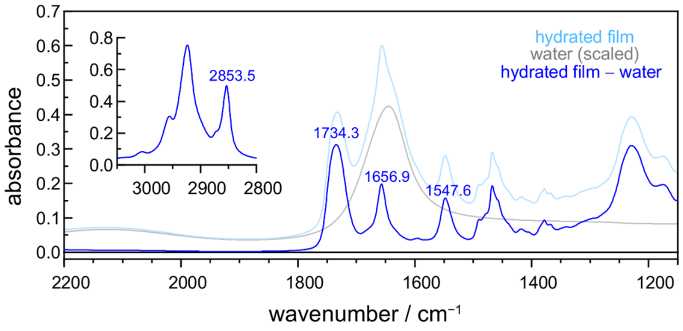


**Figure S3.** Quantifying the amount of peptide and water in the hydrated film B of KCALP-azo by FTIR absorption spectroscopy. The figure shows an unpolarized FTIR absorption spectrum of KCALP-azo reconstituted in POPC membranes (light blue trace). The contribution from water was subtracted (blue trace) using the molar absorption spectrum of $H_2O$ taken from the literature[11] (grey trace). The factor to scale the water spectrum was determined to cancel the water combination band at ~2150 $cm^{-1}$. This scaling factor, 0.019 in the present example, directly gives the amount of liquid water in $mmols/cm^2$ present in the sample.[12] Once the water absorption was subtracted, the area between 1700-1510 $cm^{-1}$ divided by 30,000 $M^{-1}cm^{-1}$ provided a rough estimate of the $mmols/cm^2$ of peptide bonds,[13] from where the $mmols/cm^2$ of peptide in the sample were derived. By this approach, we estimated 19 $\mu mol/cm^2$ of water and 20.1 $nmol/cm^2$ of peptide molecules in the depicted example (water/peptide molar ratio of ~950 / 1).

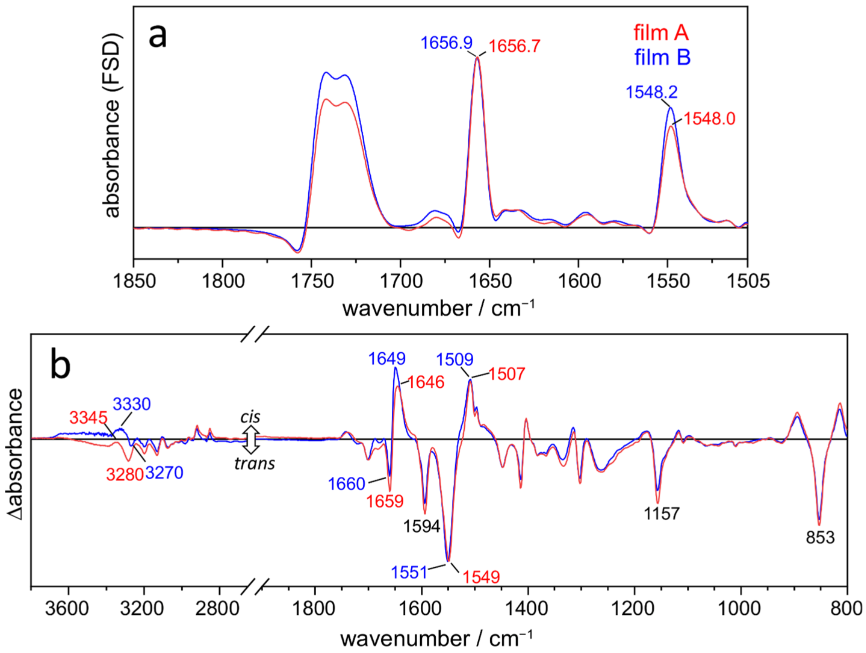


**Figure S4.** Comparison of the FTIR absorption and light-induced FTIR difference spectra of two hydrated films of KCALP-azo reconstituted in POPC bilayers: film A (red traces), with 450 water molecules per peptide, and film B (blue traces), with 950 water molecules per peptide. (a) Scaled FSD spectra ($k$ = 2 and $\gamma_L$' = 18 cm$^{-1}$), after subtraction of the water contribution. (b) Scaled light-induced FTIR difference spectra (447 nm *minus* 365 nm illumination), corresponding to the *cis-minus-trans* spectrum of KCALP-azo. Bands with maxima changing between both films are labeled. The observed shifts in the position and relative intensity of some bands might originate either from small changes in the peptide structure and light-induced structural changes as the hydration of the sample increases, or from instrumental photometric errors caused by the stronger background absorbance with increased hydration.

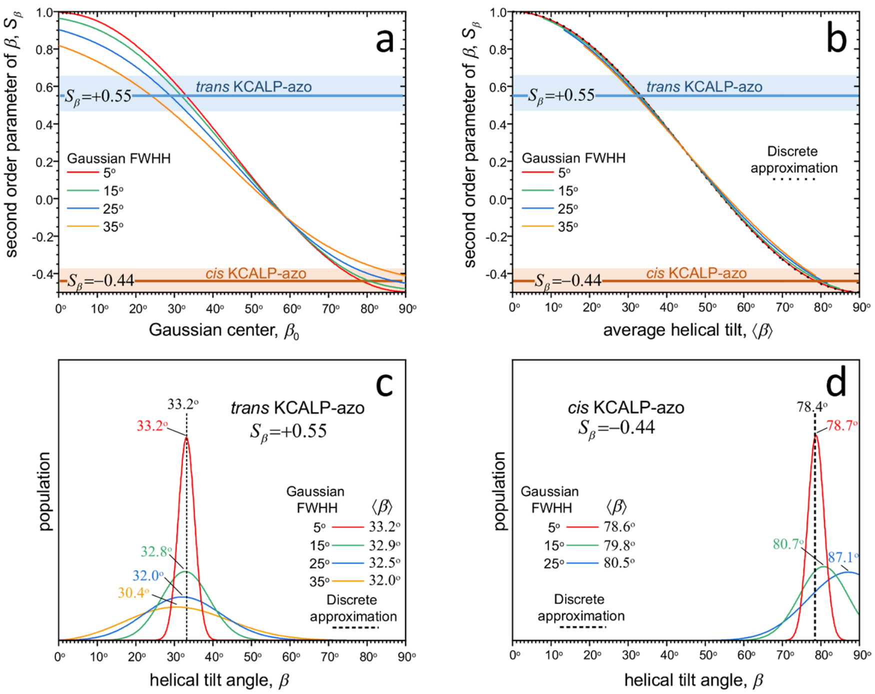


**Figure S5.** Estimation of the average helix tilt of KCALP-azo accounting for a distribution of helix tilt angles. We considered that the distributions of tilt angles follows a modified Gaussian, $Q(\beta) = N \times \exp(-4ln2(\beta - \beta_0)^2/FWHH^2)\, sin\beta$, where $\beta_0$ is the center of the Gaussian, FWHH is its full width at haft height (FWHH), $sin\beta$ is a geometric correction factor, and *N* normalizes the area of the distribution to 1 between 0º and 90º. (a) Second order parameter of the helical tilt, given by $S_\beta = (3\langle cos^2\beta\rangle - 1)/2$, as a function of $\beta_0$ for four values of FWHH (colored lines), calculated as $S_\beta = 3/2\int_0^{90} cos^2\beta \times Q(\beta)d\beta - 1/2$. (b) Second order parameter of the helical tilt as a function of the average value of the distribution of $\beta$ between 0º and 90º: $\langle\beta\rangle = \int_0^{90}\beta \times Q(\beta)d\beta$ (colored lines). The relation between $S_\beta$ and $\langle\beta\rangle$ is reproduced reasonably well using the approximation $S_\beta \approx (3cos^2\langle\beta\rangle - 1)/2$ (dotted line). (c) Several Gaussian distributions compatible with $S_\beta = +0.55$, the order parameter of the helix of *trans* KCALP-azo determined from the light-induced polarized FTIR difference spectrum (Fig. 5c). (d) Several Gaussian distributions compatible with $S_\beta = -0.44$, the order parameter of the helix of *cis* KCALP-azo determined from the light-induced polarized FTIR difference spectrum (Fig. 5c). Note how in (c) and (d) the average value of the helical tilt, $\langle\beta\rangle$, for distributions of different widths is quite similar to the value obtained using the discrete approximation. In (c) and (d) the most frequent tilt angle is indicated, whose departure from the average tilt angle increases as the distribution gets broader.

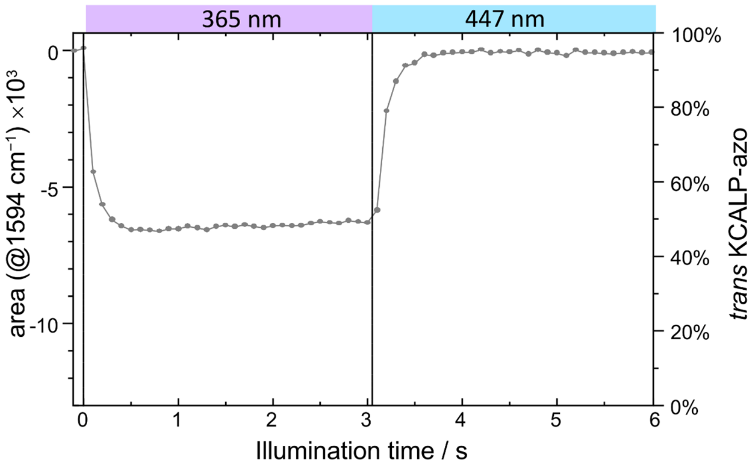


**Figure S6.** Area of the negative band at 1594 $cm^{-1}$ from the azobenzene group in the light-induced FTIR difference spectrum of KCALP-azo, as a function of the illumination time, for UV light (365 nm, 400 $mW/cm^2$) and blue light (447 nm, 200 $mW/cm^2$) illumination. To obtain the % of *trans* KCALP-azo, we assumed the photo-stationary equilibrium (PSS) reached with UV light to contain 45% of the *trans* isomer, and the PSS reached with blue light to contain 95% of the *trans* isomer (see Fig. 4a in the MS).

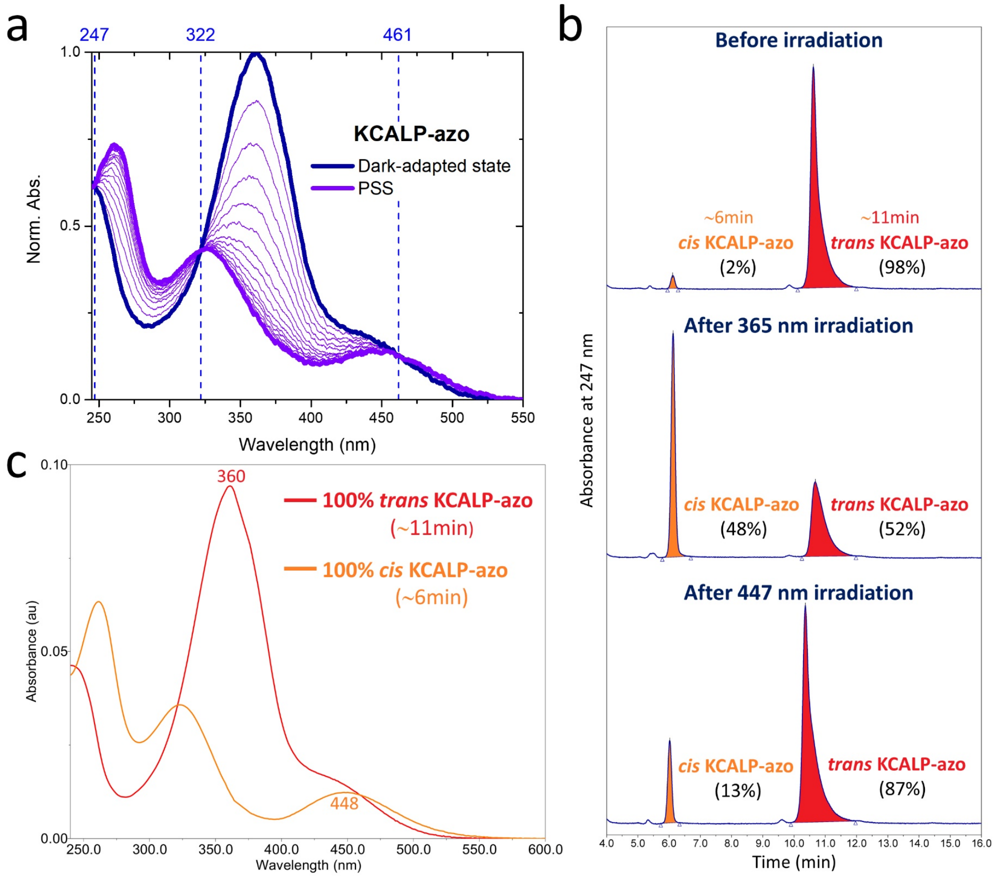


**Figure S7.** Photochromism of KCALP-azo in solution (1/1 (v/v) acetonitrile/water + 0.1% TFA). (a) UV-Vis absorption spectra of KCALP-azo under dark-adapted conditions (blue) and under continuous irradiation at 365 nm (violet), showing three isosbestic points at ~247, ~322 and ~461 nm. (b) Analytical HPLC traces of KCALP-azo with the absorbance monitored at 247 nm, before irradiation (top), after 365 nm irradiation (middle) and after 447 nm irradiation (bottom). The peak eluted at ~6 min corresponds to *cis* KCALP-azo and the peak eluted at ~11 min corresponds to *trans* KCALP-azo. By comparing the peak areas from HPLC chromatograms monitored at 247 nm, where the *cis* and *trans* isomers of KCALP-azo exhibit an equal molar absorption coefficient (see (a)), the percentage of each isomer in solution (1/1 (v/v) acetonitrile/water +0.1% TFA) was determined: 2% *cis*/98 % *trans* before irradiation, 48% *cis*/52% *trans* after 365 nm irradiation, and 13% *cis*/87% *trans* after 447 nm. (c) UV-Vis absorption spectra of the pure *cis* and *trans* isomers of KCALP-azo extracted from their corresponding elution peaks using a photodiode array detector integrated into the analytical HPLC system.

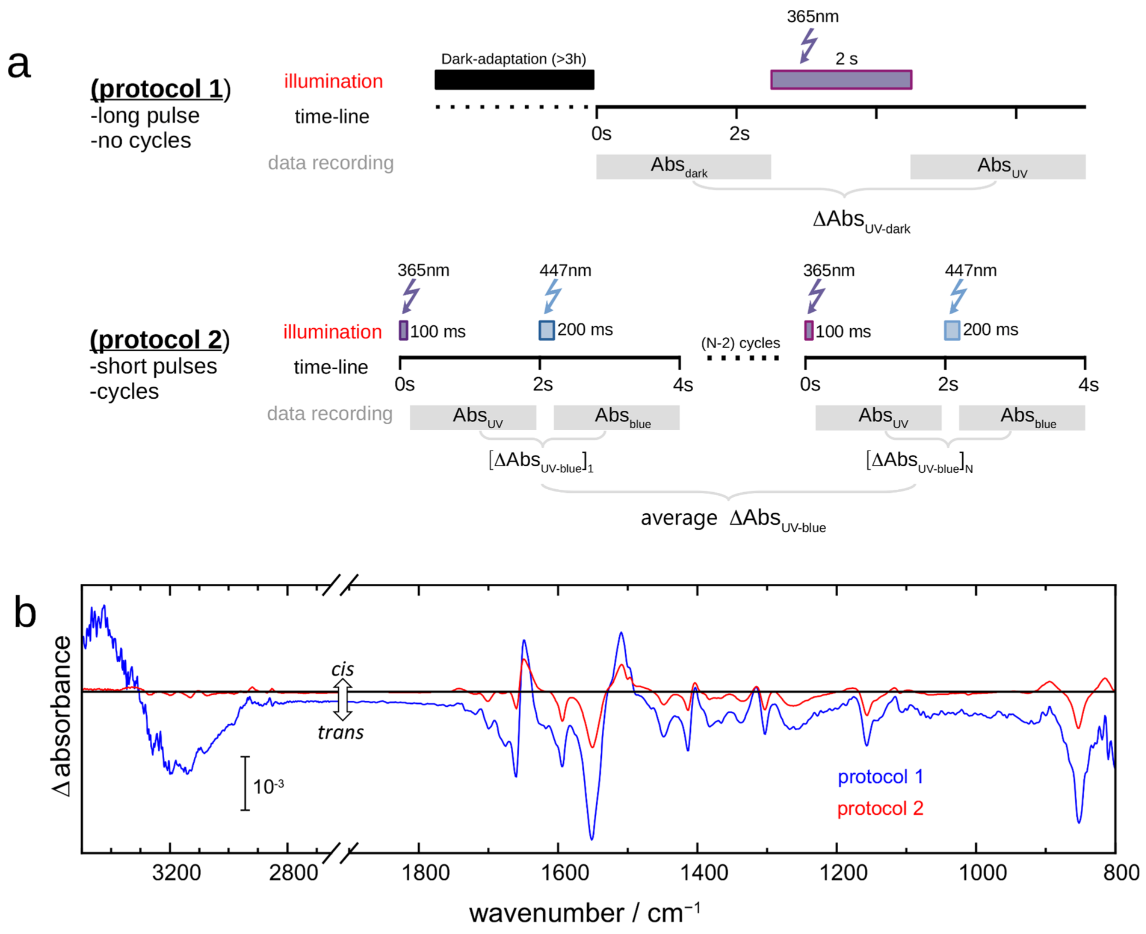


**Figure S8.** Comparison of light-induced FTIR spectra of KCALP-azo using one long UV light pulse with a single data acquisition period (protocol 1 in (a), and blue trace in (b)), versus the use of cycles alternating short UV and blue light pulses, with multiple acquisition periods (protocol 2 in (a), and red trace in (b)).

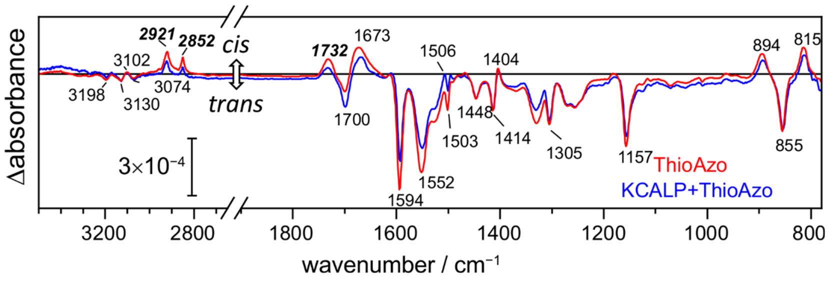


**Figure S9.** Light-induced FTIR difference spectrum of the compound ThioAzo in hydrated POPC membranes (UV-minus-blue), both in the presence (blue trace) and absence (red trace) of equimolar quantities of the peptide KCALP. The high similitude of both spectra shows that KCALP does not contribute to the measured vibrational changes under these conditions, confirming that the covalent coupling of azobenzene and KCALP is required for the latter to respond to light.

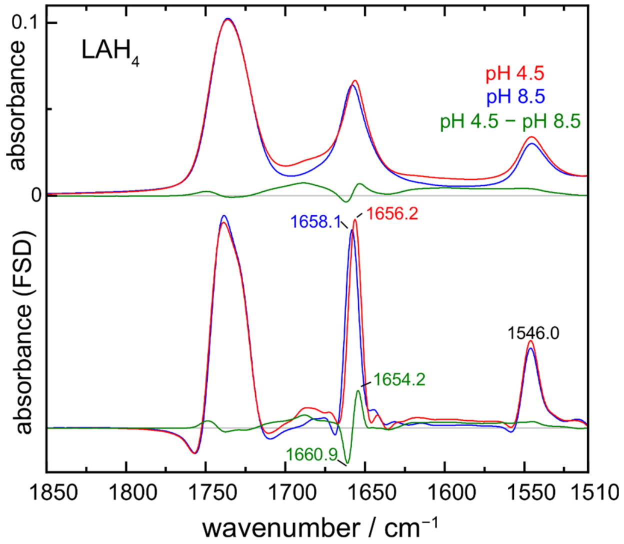


**Figure S10.** (Top) FTIR absorption spectra of the hydrophobic helical peptide $LAH_4$ in dried POPC membranes, under conditions known to adopt either a transmembrane (pH 8.5) or a membrane interface helical structure (pH 4.5).[9,14] FTIR absorption spectra measured by attenuated total reflection. Isotropic absorption spectra are shown, free from orientation effects, calculated from polarized spectra as described.[15] The pH-induced FTIR difference spectrum (pH 4.5 minus pH 8.5) is displayed in green, corresponding to the vibrational changes associated to the change in membrane topology. (Bottom) Same spectra, but after band-narrowing with FSD ($\gamma_L$' = 12 cm$^{-1}$, $k$ = 2.0). The green spectrum is reproduced in Fig. 4d of the MS.

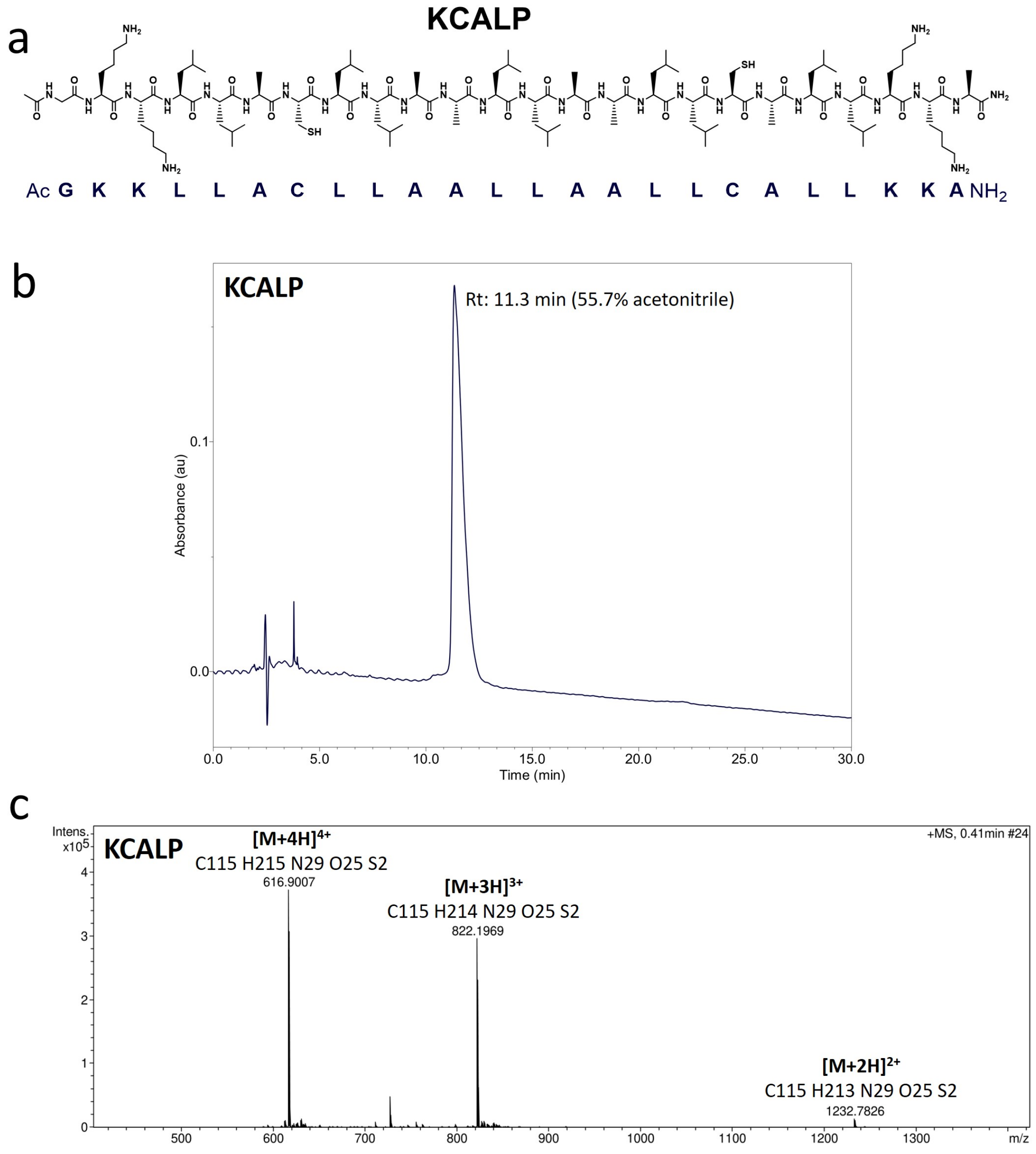


| Formula | m/z | z | \|err\| [ppm] | Mean err [ppm] | mSigma |
| --- | --- | --- | --- | --- | --- |
| C 115 H 214 N 29 O 25 S 2 | 821.8597 | 3+ | 2.6 | -3.1 | 8.7 |

**Figure S11.** Characterization of peptide KCALP. (a) Chemical structure and sequence of KCALP. (b) Analytical HPLC chromatogram of KCALP recorded at 220 nm using a linear gradient of 50-65% acetonitrile/water (+0.1 % TFA) over the course of 30 min on a Phenomenex Luna C8(2) column (5 µm, 250 mm × 4.6 mm). KCALP eluted at a retention time of 11.3 min at a flow rate of 1 mL/min, corresponding to 55.7 % acetonitrile. (c) HRMS-ESI+ spectrum of KCALP: *m/z* calc. for $C_{115}H_{214}N_{29}O_{25}S_2$ $[M+3H]^{3+}$: 821.8597, found: 822.1969.

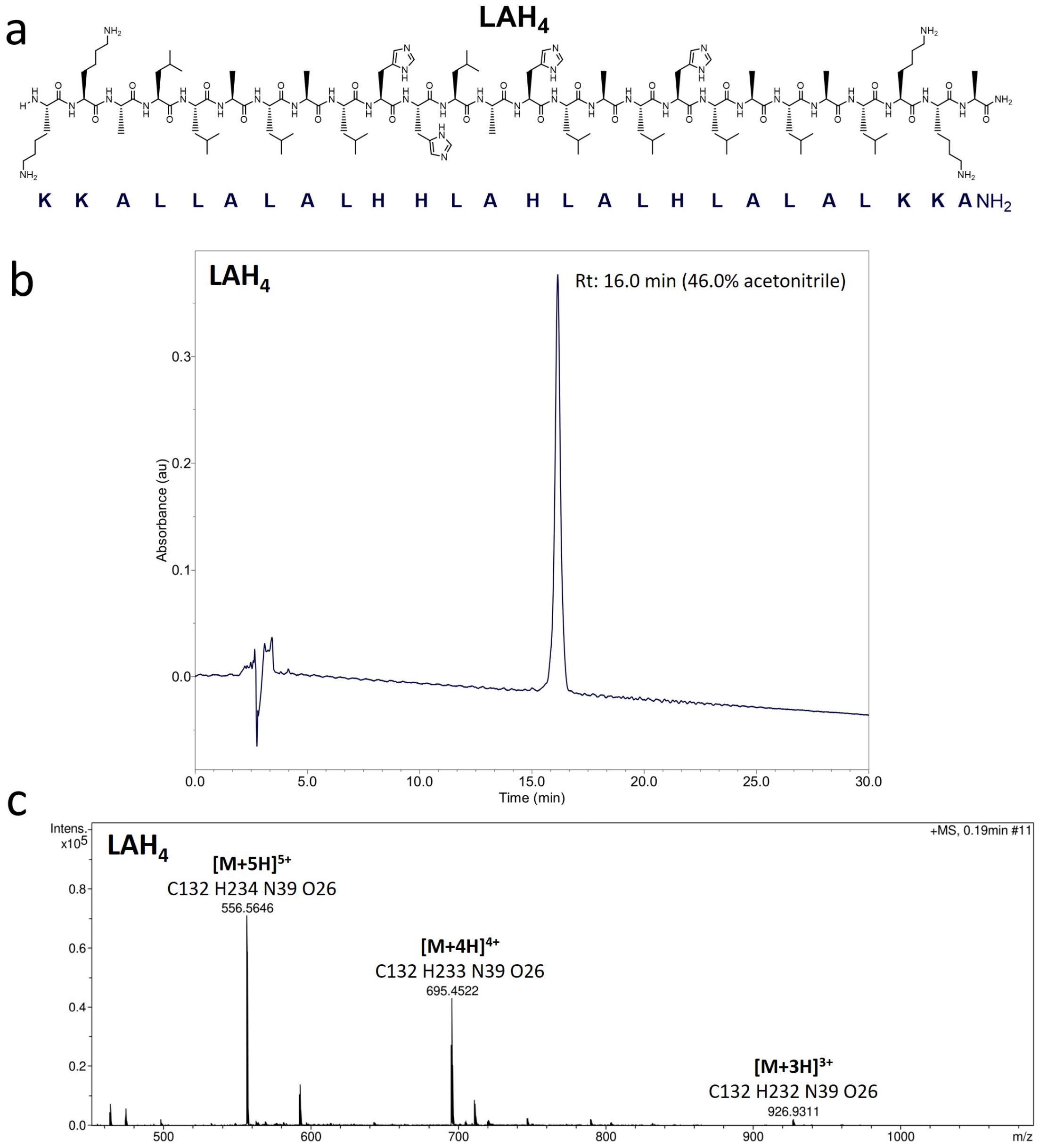


| Formula | m/z | z | \|err\| [ppm] | Mean err [ppm] | mSigma |
|---|---|---|---|---|---|
| C 132 H 233 N 39 O 26 | 695.2022 | 4+ | 0.6 | 1.6 | 8.4 |

**Figure S12.** Characterization of peptide $LAH_4$. (a) Chemical structure and sequence of $LAH_4$. (b) Analytical HPLC chromatogram of $LAH_4$ recorded at 220 nm using a linear gradient of 30-60% acetonitrile/water (+0.1 % TFA) over the course of 30 min on a Phenomenex Luna C8(2) column (5 µm, 250 mm x 4.6 mm). $LAH_4$ eluted at a retention time of 16.0 min at a flow rate of 1 mL/min, corresponding to 46.0 % acetonitrile. (c) HRMS-ESI+ spectrum of $LAH_4$: *m/z* calc. for $C_{132}H_{233}N_{39}O_{26}$ $[M+4H]^{4+}$: 695.2022, found: 695.4522.

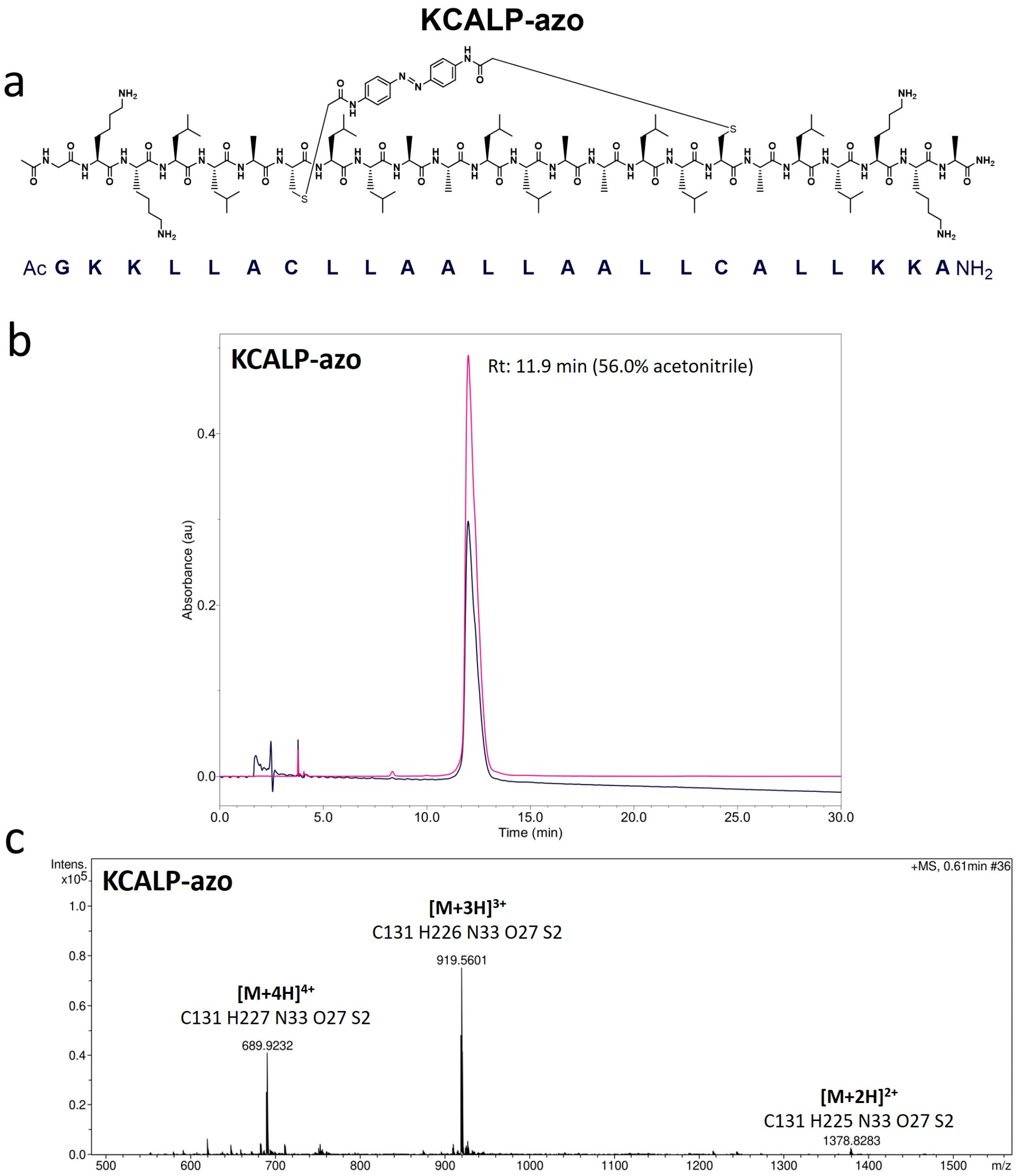


| Formula | m/z | z | \|err\| [ppm] | Mean err [ppm] | mSigma |
|---|---|---|---|---|---|
| C 131 H 226 N 33 O 27 S 2 | 919.2250 | 3+ | 0.3 | -0.2 | 7.1 |

**Figure S13.** Characterization of the photoswitchable peptide KCALP-azo. (a) Chemical structure and sequence of *trans* KCALP-azo. (b) Analytical HPLC chromatogram of KCALP-azo recorded at 220 nm (blue) and 362 nm (pink) using a linear gradient of 45-60% acetonitrile/water (+0.1 % TFA) over the course of 30 min on a Phenomenex Luna C8(2) column (5 µm, 250 mm × 4.6 mm). KCALP-azo eluted at a retention time of 11.9 min with a flow rate of 1 mL/min, corresponding to 56.0 % acetonitrile. (c) HRMS-ESI+ spectrum of KCALP-azo: *m/z* calc. for $C_{131}H_{226}N_{33}O_{27}S_2$ $[M+3H]^{3+}$: 919.2250, found: 919.5601.

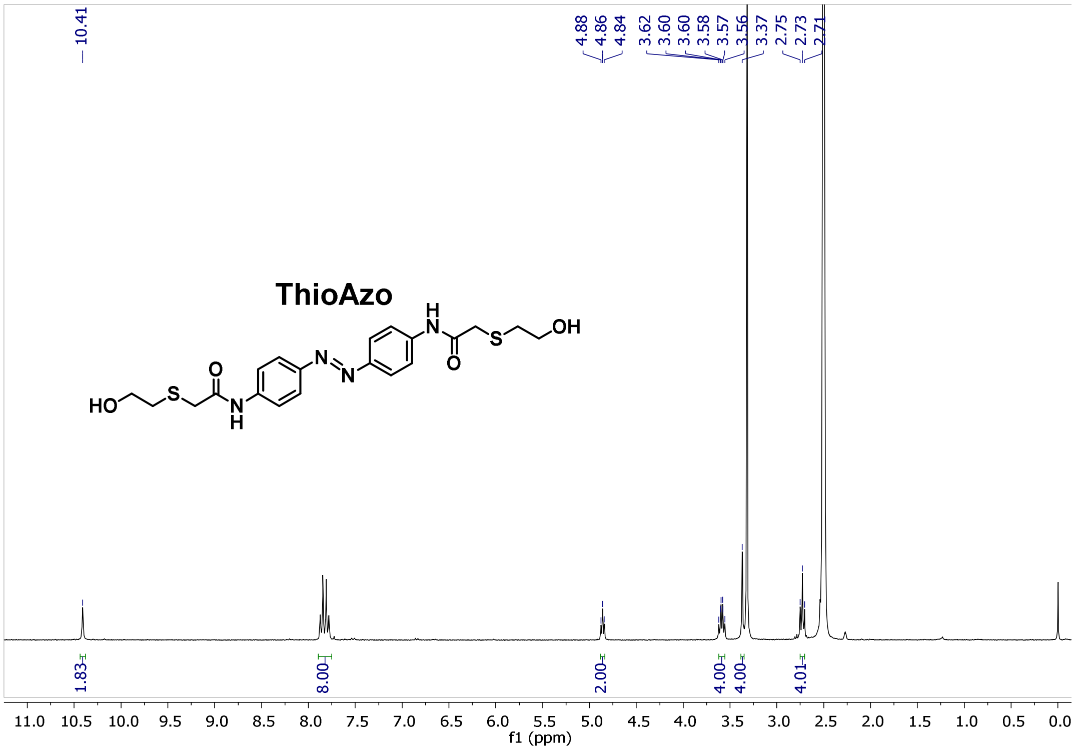


**Figure S14.** $^{1}H$ NMR spectrum of ThioAzo (300 MHz, DMSO-$d_6$). δ=10.41 (s, 2H), 7.88-7.78 (m, 8H), 4.86 (t, *J*= 5.5 Hz, 2H), 3.59 (td, *J*= 6.8, 5.4 Hz, 4H), 3.37 (s, 4H), 2.73 (t, *J*= 6.7 Hz, 4H). Coupling constants (*J*) are reported in Hz and chemical shifts values (δ) are given in parts per million (ppm), with the solvent resonance used as internal standard (DMSO-$d_6$: δ=2.50 ppm for $^{1}H$). The following abbreviations are used to indicate signal multiplicity: s (singlet), d (doublet), t (triplet) or m (multiplet).

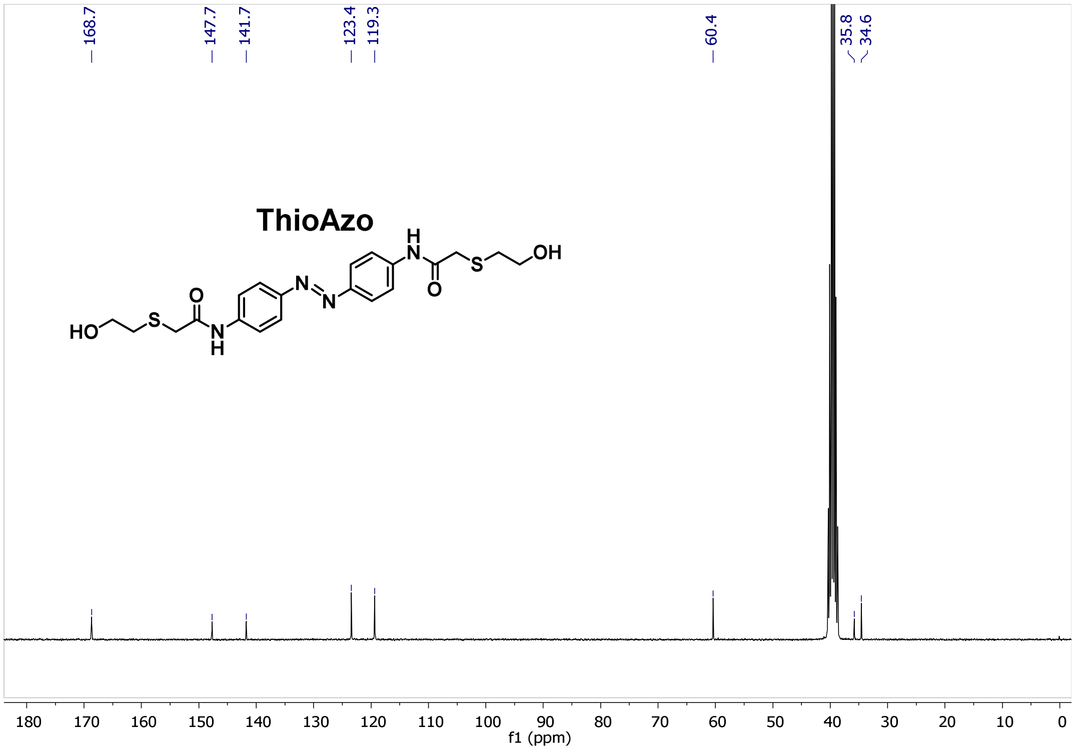


**Figure S15.** $^{13}C$ NMR spectrum of ThioAzo (75 MHz, DMSO-$d_6$). δ=168.7, 147.7, 141.7, 123.4, 119.3, 60.4, 35.8, 34.6. Chemical shifts values (δ) are given in parts per million (ppm), with the solvent resonance used as internal standard (DMSO-$d_6$: δ=39.52 ppm for $^{13}C$).

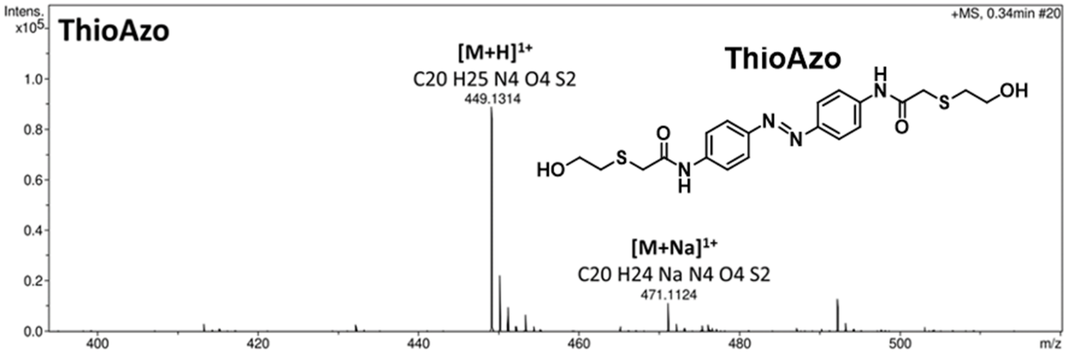


| Formula | m/z | z | \|err\| [ppm] | Mean err [ppm] | mSigma |
|---|---|---|---|---|---|
| C 20 H 25 N 4 O 4 S 2 | 449.1312 | 1+ | 0.6 | -0.7 | 9.1 |

**Figure S16.** HRMS-ESI+ spectrum of ThioAzo, with *m/z* calculated of 449.1312 and found of 449.1314 for $C_{20}H_{25}N_4O_4S_2$ $[M+H]^{1+}$.

**Supporting Tables**

**Table S1.** Band parameters and assignments of the bands in the 1850-1510 $cm^{-1}$ region of the FTIR absorption spectrum of a dry film of POPC reconstituted KCALP-azo (see Fig. 2d)

| $\nu_0$ ($cm^{-1}$) | $\gamma_L$ ($cm^{-1}$)[a] | $\gamma_G$ ($cm^{-1}$)[b] | FWHH ($cm^{-1}$) | A | A (%) | Assignment |
|---|---|---|---|---|---|---|
| **1742.2** | 4 | 15 | 18 | 3.33 | | Lipid ester $\nu$C=O |
| **1731.2** | 9 | 17 | 22 | 4.85 | | Lipid ester $\nu$C=O |
| **1696.2** | 20 | 6 | 22 | 0.47 | 8.1 | Amide I[c] |
| **1681.9** | 19 | 6 | 21 | 0.65 | 11.2 | Amide I |
| **1659.1** | 17 | 4 | 18 | 4.27 | 73.2 | Amide I (transmembrane $\alpha$-helix, $E_1$+A mode) |
| **1645.3** | 18 | 5 | 19 | 0.30 | 5.2 | Amide I |
| **1632.6** | 18 | 5 | 19 | 0.14 | 2.4 | Amide I |
| **1615.1** | 20 | 5 | 22 | 0.19 | | |
| **1597.7** | 24 | 5 | 26 | 0.72 | | C=C (azobenzene) |
| **1581.6** | 18 | 5 | 20 | 0.07 | | |
| **1546.7** | 14 | 8 | 17 | 2.14 | 68.8 | Amide II ($\alpha$-helix, $E_1$ mode) |
| **1535.4** | 14 | 10 | 19 | 0.65 | 21.0 | Amide II |
| **1520.0** | 19 | 5 | 20 | 0.32 | 10.2 | Amide II ($\alpha$-helix, A mode) |

[a] Lorentzian full-width at half height (FWHH)
[b] Gaussian FWHH
[c] from the peptide bond and from the azobenzene moiety

**Table S2.** Band parameters and assignments of the bands in the 1850-1510 cm$^{-1}$ region of the FTIR absorption spectrum of a dry film of POPC reconstituted KCALP (see Fig. 2d)

| $\nu_0$ (cm$^{-1}$) | $\gamma_L$ (cm$^{-1}$)[a] | $\gamma_G$ (cm$^{-1}$)[b] | FWHH (cm$^{-1}$) | A | A (%) | Assignment |
|---|---|---|---|---|---|---|
| **1742.2** | 5 | 15 | 18 | 3.23 | | Lipid ester $\nu$C=O |
| **1731.3** | 9 | 17 | 22 | 4.20 | | Lipid ester $\nu$C=O |
| **1695.3** | 18 | 5 | 20 | 0.19 | 2.5 | Amide I[c] |
| **1682.9** | 18 | 5 | 19 | 0.59 | 7.7 | Amide I |
| **1659.6** | 16 | 8 | 19 | 6.32 | 82.6 | Amide I (transmembrane $\alpha$-helix, $E_1$+A mode) |
| **1644.2** | 18 | 5 | 19 | 0.37 | 4.9 | Amide I |
| **1632.2** | 18 | 5 | 20 | 0.18 | 2.3 | Amide I |
| **1615.0** | 20 | 5 | 22 | 0.21 | | |
| **1599.4** | 25 | 5 | 26 | 0.51 | | |
| **1581.3** | 19 | 6 | 21 | 0.21 | | |
| **1546.8** | 13 | 7 | 17 | 3.97 | 70.7 | Amide II ($\alpha$-helix, $E_1$ mode) |
| **1536.9** | 14 | 9 | 19 | 1.21 | 21.5 | Amide II |
| **1520.4** | 19 | 6 | 21 | 0.44 | 7.8 | Amide II ($\alpha$-helix, A mode) |

[a] Lorentzian full-width at half height (FWHH)
[b] Gaussian FWHH
[c] from the peptide bond and from the azobenzene moiety

**Table S3.** Band parameters and assignments of the bands in the 1850-1510 cm$^{-1}$ region of the FTIR absorption spectrum of a hydrated film of POPC reconstituted KCALP-azo (film A, 450 water molecules / peptide, see Fig. 3c).

| $\nu_0$ (cm$^{-1}$) | $\gamma_L$ (cm$^{-1}$)[a] | $\gamma_G$ (cm$^{-1}$)[b] | FWHH (cm$^{-1}$) | A | A (%) | Assignment |
|---|---|---|---|---|---|---|
| **1744.6** | 2 | 15 | 16 | 1.56 | | Lipid ester νC=O |
| **1731.9** | 3 | 21 | 23 | 3.31 | | Lipid ester νC=O |
| **1719.8** | 18 | 18 | 29 | 2.51 | | Lipid ester νC=O |
| **1679.9** | 20 | 6 | 22 | 0.44 | 8.8 | Amide I |
| **1656.8** | 15 | 4 | 16 | 3.26 | 65.9 | Amide I (transmembrane α-helix, $E_1$+A mode) |
| **1646.0** | 19 | 6 | 21 | 0.64 | 12.8 | Amide I (hydrated α-helix, $E_1$+A mode) |
| **1633.2** | 20 | 7 | 22 | 0.61 | 12.4 | Amide I |
| **1615.5** | 19 | 6 | 20 | 0.16 | | |
| **1595.7** | 20 | 6 | 22 | 0.47 | | C=C (azobenzene) |
| **1578.0** | 20 | 6 | 22 | 0.21 | | |
| **1547.8** | 14 | 9 | 19 | 2.34 | 70.7 | Amide II (α-helix, $E_1$ mode) |
| **1535.3** | 15 | 10 | 20 | 0.70 | 21.2 | Amide II |
| **1518.7** | 19 | 6 | 20 | 0.27 | 8.0 | Amide II (α-helix, A mode) |

[a] Lorentzian full-width at half height (FWHH)
[b] Gaussian FWHH

**Table S4.** Band parameters and assignments of the bands in the 1850-1510 cm$^{-1}$ region of the FTIR absorption spectrum of a hydrated film of POPC reconstituted KCALP-azo (film B, 950 water molecules / peptide, see Fig. S3).

| $\nu_0$ (cm$^{-1}$) | $\gamma_L$ (cm$^{-1}$)[a] | $\gamma_G$ (cm$^{-1}$)[b] | FWHH (cm$^{-1}$) | A | A (%) | Assignment |
|---|---|---|---|---|---|---|
| **1745.2** | 5 | 14 | 17 | 2.35 | | Lipid ester $\nu$C=O |
| **1733.3** | 4 | 22 | 24 | 5.66 | | Lipid ester $\nu$C=O |
| **1722.2** | 22 | 15 | 30 | 5.05 | | Lipid ester $\nu$C=O |
| **1679.7** | 25 | 9 | 28 | 1.29 | 17.3 | Amide I |
| **1657.0** | 16 | 4 | 17 | 4.62 | 61.8 | Amide I (transmembrane $\alpha$-helix, $E_1$+A mode) |
| **1645.6** | 19 | 6 | 21 | 0.58 | 7.8 | Amide I (hydrated $\alpha$-helix, $E_1$+A mode) |
| **1633.1** | 22 | 8 | 25 | 0.98 | 13.1 | Amide I |
| **1616.4** | 20 | 6 | 22 | 0.33 | | |
| **1596.2** | 20 | 6 | 22 | 0.69 | | C=C (azobenzene) |
| **1577.4** | 23 | 9 | 26 | 0.43 | | |
| **1548.3** | 14 | 8 | 18 | 3.59 | 70.6 | Amide II ($\alpha$-helix, $E_1$ mode) |
| **1537.0** | 15 | 10 | 20 | 1.08 | 21.1 | Amide II |
| **1519.6** | 20 | 7 | 23 | 0.42 | 8.3 | Amide II ($\alpha$-helix, A mode) |

[a] Lorentzian full-width at half height (FWHH)
[b] Gaussian FWHH

**Table S5.** Band parameters and assignments of the bands in the light-induced FTIR difference spectrum of KCALP-azo of a hydrated film of POPC reconstituted KCALP-azo (film B, 950 water molecules / peptide, see Fig. 4e). Positive and negative bands correspond to the *cis* and *trans* conformations of KCALP-azo, respectively.

| $\nu_0$ (cm$^{-1}$) | $\gamma_L$ (cm$^{-1}$)[a] | $\gamma_G$ (cm$^{-1}$)[b] | FWHH (cm$^{-1}$) | A×10$^3$ | A (%) | Assignment |
|---|---|---|---|---|---|---|
| **1742.6** | 0 | 16 | 16 | 1.3 | | Lipid ester νC=O |
| **1721.0** | 0 | 12 | 12 | 0.5 | | Lipid ester νC=O (*trans*) |
| **1701.9** | 0 | 11 | 11 | −2.2 | | |
| **1688.8** | 0 | 9 | 9 | 0.4 | | |
| **1680.3** | 10 | 6 | 13 | −0.7 | −0.9 | Amide I peptide (*trans*) |
| **1668.1** | 12 | 5 | 14 | 3.4 | 3.8 | Amide I peptide (*cis*) |
| **1657.2** | 13 | 4 | 14 | −77.5 | −99.1 | Amide I peptide (*trans*) |
| **1655.6** | 13 | 5 | 15 | 76.4 | 86.2 | Amide I peptide (*cis*) |
| **1643.5** | 13 | 8 | 17 | 8.8 | 10.0 | Amide I peptide (*cis*) |
| **1608.8** | 9 | 4 | 11 | 1.8 | | |
| **1605.4** | 9 | 4 | 11 | −2.7 | | |
| **1594.7** | 4 | 6 | 9 | −6.4 | | C=C photoswitch (*trans*) |

[a] Lorentzian full-width at half height (FWHH)
[b] Gaussian FWHH

**Supplementary References**


(1) Kimura, Y.; Vassylyev, D. G.; Miyazawa, A.; Kidera, A.; Matsushima, M.; Mitsuoka, K.; Murata, K.; Hirai, T.; Fujiyoshi, Y. Surface of Bacteriorhodopsin Revealed by High-Resolution Electron Crystallography. *Nature* **1997**, *389* (6647), 206–211. https://doi.org/10.1038/38323.

(2) Pebay-Peyroula, E.; Dahout-Gonzalez, C.; Kahn, R.; Trézéguet, V.; Lauquin, G. J. M.; Brandolin, G. Structure of Mitochondrial ADP/ATP Carrier in Complex with Carboxyatractyloside. *Nature* **2003**, *426* (6962), 39–44. https://doi.org/10.1038/nature02056.

(3) Ethayathulla, A. S.; Yousef, M. S.; Amin, A.; Leblanc, G.; Kaback, H. R.; Guan, L. Structure-Based Mechanism for $Na^+$/Melibiose Symport by MelB. *Nat. Commun.* **2014**, *5*, 3009. https://doi.org/10.1038/ncomms4009.

(4) Lórenz-Fonfría, V. A.; León, X.; Padrós, E. Studying Substrate Binding to Reconstituted Secondary Transporters by Attenuated Total Reflection Infrared Difference Spectroscopy. *Methods Mol. Biol.* **2012**, *914*, 107–126. https://doi.org/10.1007/978-1-62703-023-6_7.

(5) Lórenz, V. A.; Villaverde, J.; Trézéguet, V.; Lauquin, G. J.-M.; Brandolin, G.; Padrós, E. The Secondary Structure of the Inhibited Mitochondrial ADP/ATP Transporter from Yeast Analyzed by FTIR Spectroscopy. *Biochemistry* **2001**, *40* (30), 8821–8833. https://doi.org/10.1021/bi010091s.

(6) Lorenz-Fonfria, V. A.; Saita, M.; Lazarova, T.; Schlesinger, R.; Heberle, J. PH-Sensitive Vibrational Probe Reveals a Cytoplasmic Protonated Cluster in Bacteriorhodopsin. *Proc. Natl. Acad. Sci. U. S. A.* **2017**, *114* (51), E10909–E10918. https://doi.org/10.1073/pnas.1707993114.

(7) Kabsch, W.; Sander, C. Dictionary of Protein Secondary Structure: Pattern Recognition of Hydrogen-bonded and Geometrical Features. *Biopolymers* **1983**, *22* (12), 2577–2637. https://doi.org/10.1002/bip.360221211.

(8) Klose, D. P.; Wallace, B. A.; Janes, R. W. 2Struc: The Secondary Structure Server. *Bioinformatics* **2010**, *26* (20), 2624–2625. https://doi.org/10.1093/bioinformatics/btq480.

(9) Bechinger, B. Towards Membrane Protein Design: PH-Sensitive Topology of Histidine-Containing Polypeptides. *J. Mol. Biol.* **1996**, *263* (5), 768–775. https://doi.org/10.1006/JMBI.1996.0614.

(10) Goormaghtigh, E.; Raussens, V.; Ruysschaert, J.-M. Attenuated Total Reflection Infrared Spectroscopy of Proteins and Lipids in Biological Membranes. *Biochim. Biophys. Acta - Rev. Biomembr.* **1999**, *1422* (2), 105–185. https://doi.org/10.1016/S0304-4157(99)00004-0.

(11) Bertie, J. E.; Lan, Z. Infrared Intensities of Liquids XX: The Intensity of the OH Stretching Band of Liquid Water Revisited, and the Best Current Values of the Optical Constants of $H_2$ O(l) at 25°C between 15,000 and 1 $Cm^{-1}$. *Appl. Spectrosc.* **1996**, *50* (8), 1047–1057. https://doi.org/10.1366/0003702963905385.

(12) Lorenz-Fonfria, V. A. Infrared Difference Spectroscopy of Proteins: From Bands to Bonds. *Chem. Rev.* **2020**, *120* (7), 3466–3576. https://doi.org/10.1021/acs.chemrev.9b00449.

(13) Rahmelow, K.; Hübner, W.; Ackermann, T. Infrared Absorbances of Protein Side Chains. *Anal.*

*Biochem.* **1998**, *257* (1), 1–11. https://doi.org/10.1006/abio.1997.2502.

(14) Bechinger, B.; Ruysschaert, J.-M.; Goormaghtigh, E. Membrane Helix Orientation from Linear Dichroism of Infrared Attenuated Total Reflection Spectra. *Biophys. J.* **1999**, *76* (1), 552–563. https://doi.org/10.1016/S0006-3495(99)77223-1.

(15) Lórenz-Fonfría, V. A.; Granell, M.; León, X.; Leblanc, G.; Padrós, E. In-Plane and Out-of-Plane Infrared Difference Spectroscopy Unravels Tilting of Helices and Structural Changes in a Membrane Protein upon Substrate Binding. *J. Am. Chem. Soc.* **2009**, *131* (42), 15094–15095. https://doi.org/10.1021/ja906324z.
